# Nutrient Source and Mycorrhizal Association jointly alters Soil Microbial Communities that shape Plant-Rhizosphere-Soil Carbon-Nutrient Flows

**DOI:** 10.1101/2020.05.08.085407

**Authors:** Somak Chowdhury, Markus Lange, Ashish A Malik, Timothy Goodall, Jianbei Huang, Robert I Griffiths, Gerd Gleixner

**Author notes:** Corresponding author: Gerd Gleixner, Institutional Affiliation: Department of Biogeochemical Processes, Max Planck Institute for Biogeochemistry, 07745, Jena, Germany.

## Abstract

Interactions between plants and microorganisms strongly affect ecosystem functioning as processes of plant productivity, litter decomposition and nutrient cycling are controlled by both organisms. Though two-sided interactions between plants and microorganisms and between microorganisms and litter decomposition are areas of major scientific research, our understanding of the three-sided interactions of plant-derived carbon flow into the soil microbial community and their follow-on effects on ecosystem processes like litter decomposition and plant nutrient uptake remains limited. Therefore, we performed a greenhouse experiment with two plant communities differing in their ability to associate with arbuscular mycorrhizal fungi (AMF). By applying a ^13^CO_2_ pulse label to the plant communities and adding various ^15^N labelled substrate types to ingrowth cores, we simultaneously traced the flow of plant-derived carbon into soil microbial communities and the return of mineralized nitrogen back to the plant communities. We observed that net ^13^C assimilation by the rhizosphere microbial communities and their community composition not only depended on plant-AMF association but also type of substrate being decomposed. AMF-association resulted in lower net ^13^C investment into the decomposer community than absence of the association for similar ^15^N uptake. This effect was driven by a reduced carbon flow to fungal and bacterial saprotrophs and a simultaneous increase of carbon flow to AMF. Additionally, in presence of AMF association CN flux also depended on the type of substrate being decomposed. Lower net ^13^C assimilation was observed for decomposition of plant-derived and microorganism-derived substrates whereas opposite was true for inorganic nitrogen. Interestingly, the decomposer communities assembled in the rhizosphere were structured by both the plant community and substrate amendments which suggests existence of functional overlap between the two soil contexts. Moreover, we present preliminary evidence that AMF association helps plants access nutrients that are locked in bacterial and plant necromass at a lower carbon cost. Therefore, we conclude that a better understanding of ecosystem processes like decomposition can only be achieved when the whole plant-microorganism-litter context is investigated.

## Introduction

Plant growth is mostly limited by nutrient availability in soil, as nutrients in soil are either soil organic matter bound or associated with the mineral matrix (Veresoglou et al., 2012). Consequently, plants invest substantial parts of their photoassimilated carbon into soil via root exudates (sugars, amino acids and organic acids) and rhizodeposits, to stimulate microbial decomposition of soil organic matter and nutrient release (Jacoby et al., 2017; Kuzyakov and Xu, 2013). Additionally, root exudates can also alter organo-mineral associations (Keiluweit et al., 2015), pH (Malik et al., 2018) and redox potential (Husson, 2013) of the soil to directly improve nutrient availability. However, immediate plant uptake of available nutrients is not always possible (Dijkstra et al., 2013) since soil microbial communities have a high nutrient demand and often assimilate nutrients more efficiently than plants (Levin and Angert, 2015; Zhu et al., 2017). Since nutrient cycles in soil are largely controlled by the various biotic (root and microbial traits) and abiotic soil properties (minerology, pH and redox), variations in root exudation patterns and association between plants and microorganisms can have a tremendous impact on the rates of nutrient cycling in the rhizosphere (Craine et al., 2007; Veresoglou et al., 2012).

To enhance nutrient uptake from soil a majority of terrestrial plants species (> 80%) rely on direct root association with arbuscular mycorrhizal fungi (AMF) (Bonfante and Genre, 2010; Davison et al., 2015). Though AMF can’t decompose organic matter themselves (Tisserant et al., 2013), they can use their hyphal network to scavenge nutrients and compete with the decomposer community for nutrient uptake (Averill et al., 2019; Bunn et al., 2019). The AMF network also benefits plants by allowing indirect root access to a greater soil volume (Sanders and Tinker, 1973), immobile nutrient pools (Hestrin et al., 2019) and distant water pockets (Bowles et al., 2018; Karlowsky et al., 2018), thus greatly increasing the area across which plant carbon is distributed. As a result, presence of AMF association ensures that fresh photoassimilates are allocated to the soil microbial community in a targeted manner through the hyphal network (Kaiser et al., 2015) for nutrient exchange. The extent of this exchange however is regulated by the availability of nutrients in soil (Whiteside et al., 2019). In contrast, an absence of AMF association results in passive release of photoassimilated carbon across comparatively reduced soil volumes (Kaiser et al., 2015). To understand how these distinct modes of carbon delivery into soil affect nutrient cycles and plant microbial nutrient fluxes, it is essential to examine them in the context of various nutrient sources from which plants generally acquire nutrients.

Microbial decomposition processes are critical for sustained nutrient release in soil (Schimel and Bennett, 2004), however they are limited by the availability of labile carbon (Hütsch et al., 2002) - a primary source of which are root exudates. The same root exudates can also destabilize mineral nutrient associations and ensure nutrient release for the decomposer community (Jilling et al., 2018). As root-AMF associations can alter both the quantity (de Graaff et al., 2010) and biochemical profile (Shi et al., 2016; Zhalnina et al., 2018) of root exudates, they can have a large impact on nutrient availability in soil. Similarly, nutrient content and type of decomposing organic matter can significantly alter associated decomposer communities and their activity (Güsewell and Gessner, 2009; Meier et al., 2015; Strickland et al., 2009) to influence rates of nutrient cycling in soil. Recent findings based on functional analysis of rhizosphere microbial communities strongly indicate that a functionally diverse microbial community is maintained for exudate utilization in the rhizosphere compared to the bulk soil (Nuccio et al., 2020). Though exudate and litter associated shifts in the microbial community of the rhizosphere are well known, understanding the underlying carbon-nutrient flux between plant-AMF association and these decomposer communities across various nutrient sources can offer further insights into mechanisms of plant nutrient uptake.

Soil receives nitrogen containing substances either via carbon rich plant residues, nitrogen rich microbial residues or inorganic nitrogen from deposition and fertilizers (Gleixner, 2013; Liang et al., 2017; Thuille et al., 2015). However, the difference amongst these inputs based on nutrient content (litter quality) often hides the effect of their biochemical characteristics (litter type) on decomposer community assembly and nutrient exchange. Under natural conditions, the decomposer community is typically shaped by litter characteristics and their physiological boundaries, defined mainly by their energetic and nutritional needs (Thomson et al., 2013). As organic matter decomposition is mostly mediated by microbial communities, a sizable portion of the soil carbon and nitrogen pool is of microbial origin (Xu et al., 2013). Moreover, the porous nature of soil matrix and presence of complex surfaces ensures that nutrient rich microbial residues are bound within nutrient poor exopolysaccharides as part of soil biofilms (Kallenbach et al., 2016). Though exchange of nutrients between the microbial pool and plants are commonplace, however, unlike plant root litter, little is known about how their turnover impacts carbon-nutrient flux during litter decomposition in the rhizosphere (Nuccio et al., 2020), thus underscoring the need for further investigation through labelled nutrient amendments.

In summary, large uncertainties still exist regarding the mechanistic benefits of plant-AMF associations with respect to organic matter source and nutrient return. Therefore, we assembled two plant communities that differed in their ability to associate with AMF (AM or NM) and amended the root colonized soil with three ^15^N labeled nitrogen substrates (plant root litter, microbial necromass and inorganic N) of similar quality (C:N ratio) and traced the net plant derived carbon flow into the microbial community (PLFA/NLFA) using a ^13^CO_2_ pulse label experiment. We also investigated the microbial community composition using DNA and RNA based amplicon sequencing. We first hypothesized that AMF associations result in greater plant carbon export to the microbial community (including AMF) for nitrogen mineralization. We also hypothesized that carbon cost per nutrient gain is higher in presence of AMF association than absence, since AMF associations require considerable amounts of plant carbon for maintaining the hyphal network. We finally hypothesize that the associated soil microbial community depends in the first order on the plant community (AMF vs Non-AMF) and its carbon export, and in the second order on the substrate origin independent from the total amount of added nutrients.

## Materials and Methods

### Microcosm Experiment

The soil used for the experiment was collected from the Jena Experiment - a grassland biodiversity experiment located in Jena, Germany in March 2015. The soil type was Eutric Fluvisol (Roscher et al., 2004) which has a pH of 7.4 (s.d. 0.06), organic carbon content of 21.1 g C kg^-1^ (s.d. 3.6 g C kg^-1^) and total nitrogen content of 2.2 g C kg^-1^ (s.d. 0.3 g N kg^-1^) (Weisser et al., 2017). The soil contains a diverse mixture of bacteria and fungi (Weisser et al., 2017) that served as a natural inoculum in our experiment. After removing the root mat in the upper 2 cm, the top 5 cm of soil was collected, sieved (<2 mm) and all remaining roots were removed. The soil was well mixed and stored at 4 °C until the establishment of the pots 1 week later. We set up microcosms (19 × 12.5 × 9.5 cm) and established mixed temperate plant communities of two types namely AMF dependent plant community (*Linum ussitatissimum, Linum perenne, Festuca rubra* and *Tagetes patula*) termed as the AM plant community and plant community lacking in AMF associations (*Dianthus caryophyllus, Matthiola longipetala sub-species bicornis, Dianthus barbaratus* and *Carex arenaria*) termed as NM plant community (Brundrett, 2015b, 2015a). By using plant communities instead of monocultures, we intended to avoid well-known species-specific effects of root exudation. Therefore we used 4 species mixtures that have already been shown to overcome species specific effects (Weisser et al., 2017). Therefore, our experiment provides plant community level understanding of nutrient flux between plants and microorganisms.

Each microcosm had a single central PVC ingrowth core (diameter = 5 cm) with two tiers of 8 windows (4 × 1.5 cm) guarded by a 1000 micron mesh (Figure S1). No external inoculum of AMF was used as the source soil already contained a diverse AMF community as confirmed by types of plants prevalent at the site (Harley and Harley, 1987). For both plant communities (AM and NM) we used a 1000 micron mesh to ensure the growth of fine roots and hyphae into the ingrowth core (Figure S1). We confirmed root access at the time of substrate amendment and final harvest (Figure S2). The following replication scheme was implemented in the experiment 3 biological replicates x 3 soil amendments x 4 timepoints resulting in a total of 36 independent microcosms per plant community. The microcosms and the ingrowth cores were filled with well mixed soil at field bulk density and were pre-equilibrated for 2 weeks in the greenhouse. Even plant communities for each plant community type were established by planting 1 seedling per species that already developed first true leaves within each microcosm unit outside the ingrowth core resulting in a final plant species richness of 4 in each microcosm (Figure 1). The microcosms were randomly distributed in the greenhouse and maintained for 4 months until the plants were fully developed. Irrigation was provided using an ebb and flood system for 5 min daily and lighting for the plants (Son-T Agro 430 W HPS bulbs, primary light range = 520–610 nm, Philips Lighting Company, Somerset, NJ, USA) was provided daily for 12 h. Plant species richness and evenness were visually monitored throughout the experiment until the time of harvest.

**Figure 1:**
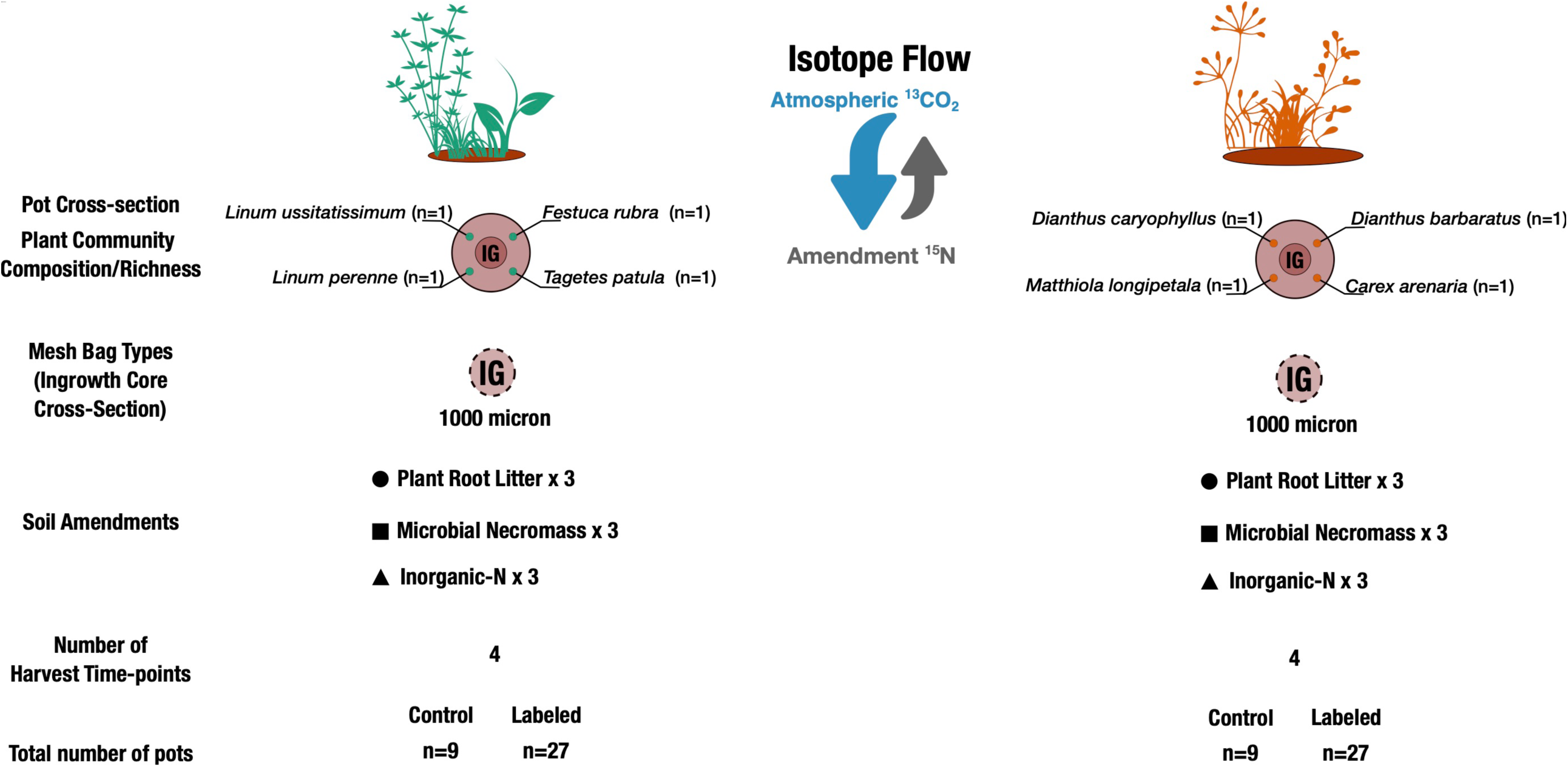
Experimental setup of the study.

In order to make substrate amendments to fully developed microcosms, the central ingrowth core was removed. The soil was sieved (<2mm) to remove fine roots (Refer Section 1.2 of Supplementary Information). The N amendments made to ingrowth core soil differed in their origin (plant, microbial and inorganic). The plant root litter and microbial necromass differed in their native C:N ratio with plant root litter having a C:N ratio of 45.67 ± 1.09 (C content = 7.41 % and N content = 0.16 %) and microbial necromass had a C:N ratio of 8.37 ± 1.35 (C content = 12.88 % ; N content = 1.54 %). For plant root litter amendment, sieved ingrowth core soil was uniformly mixed with 7.5 g of shredded plant root litter which resulted in an input of 48.55 μg of ^15^N into the soil. For microbial necromass amendments, sieved soil was amended with 0.8 g of microbial necromass containing 49.53 μg of ^15^N with an additional 1.28 g of nitrogen free xanthan resulting in an additional carbon amendment of 0.448 g which ensured a similar input of carbon and nitrogen as that of plant root litter. The inorganic-N amendment with KNO_3_ was carried out by mixing 0.012 g of KNO_3_ (^15^N content = 48.25 μg) with 1.58 g of xanthan resulting in a total C input of 0.553 g C, thus ensuring a similar carbon and nitrogen input as that of plant root litter amendment. Overall, the amendments were kept small, the total carbon and nitrogen content of the ingrowth core soil increased by ca. 0.15 % and ca. 0.01 %, respectively. The amended soil was mixed, refilled into the ingrowth core and reinstalled on the microcosm.

2 weeks after the amendment, which allowed roots and hyphae to regrow into the ingrowth core, the microcosms were exposed to a ^13^CO_2_ pulse in an air tight plexiglass chamber (Malik et al., 2015). For each litter amendment, all microcosms (3 replicates x 2 plant treatments x 3 post-labelling timepoints = 18 microcosms) were subjected to a ^13^CO_2_ pulse together in a well-lit (PAR intensity >500 µmol photons m^-2^s^-1^) chamber (volume = 0.15 m^3^). The chamber air was flushed with CO_2_ free synthetic air until the CO_2_ concentration fell below 150 ppm. Enriched ^13^CO_2_ (99 atom %) was introduced into the chamber using a mass flow controller (model-MF1-20sccm-N2, MKS Instruments Inc.) at a flow rate of 3.33 × 10^−7^ m^3^s^-1^ for 10 min and circulated using internal fans. The CO_2_ concentration was monitored online by a cavity ringdown spectrometer (G2101-I, Picarro Inc., Santa Clara, CA, USA) throughout the labeling period of 1h. An additional maintenance stream of ^13^CO_2_ was introduced halfway at a flow rate of 1.67 × 10^−7^ m^3^s^-1^ for 5 min that kept the chamber CO_2_ concentration between 350 ppm and 410 ppm. The chamber temperature was maintained close to greenhouse temperature by placing racks of frozen ice packs along the vertical walls of the chamber and replaced for every batch of pots that were labelled.

At the end of the labelling period the microcosms were removed from the chamber and returned back to the greenhouse and destructively harvested at 4 h, 24 h and 240 h from the pulse. The soil from ingrowth core was completely inhabited by roots (Figure S2) suggesting that all soil in the ingrowth core was rhizosphere soil. The roots were removed by sieving to 2mm sieve and hand picking (SI Section 1.2) and remaining rhizosphere soil was divided into three sub-samples and stored at -80 °C for molecular analysis within 1 h of harvest, dried at 105 °C for 48 h and stored at room temperature for elemental analysis and at -20 °C for microbial biomass and respiration analysis. The above and below ground plant tissues were collected at the last destructive harvest (240 h) and dried at 60 °C for 48 h and subsequently stored for future analysis. Prior to ^13^CO_2_ labelling of microcosms, control pots for each litter amendment (3 replicates x 3 soil amendments x 2 plant community treatments = 18 microcosms) were also placed for an hour in the chambers with similar synthetic CO_2_ free air and ^12^CO_2_ pulsing regimes as the labelled samples with similar temperature maintenance and chamber ventilation. To avoid cross contamination from ^13^C labeled air, the control microcosms were harvested prior to labelled pots.

### Production of labelled substrates

To follow nutrients released by decomposition to the plant tissues, ^15^N labelled organic matter of three types namely plant root litter, microbial necromass and synthetic substrate were used in this study. Plant root litter was harvested from perennial grass *Holcus lanatus* grown in a pot (57x 37 × 20 cm) containing sand in a plant growth chamber that was lit daily for 12 h at day temperatures of 23 **°**C and at 63 % relative humidity. The plants were watered 2-3 times a week and fertilized weekly with a balanced N free Hoagland’s solution amended with 0.40 atom% K^15^NO_3_ as the sole nitrogen source. At the end of 4 weeks the plants were harvested and roots were cleaned with distilled water, dried and shredded to remove effects of particle size.

Inoculum for producing microbial necromass was generated by slow centrifugation of 6 g of Jena Experiment soil in 20 ml of distilled water at 650 g for 15 min. The supernatant was inoculated into M9 minimal medium amended with 20% glucose and contained K^15^NO_3_ (0.40 atom%) as only nitrogen source. The culture grew for 7 days at 19 **°**C on a horizontal shaker. The entire cell mass was harvested by centrifugation (@ 3500 g for 15 min) and re-suspended in 10 ml of sterile distilled water and sterilized by autoclaving. The sterile pellet was freeze dried for further use.

The inorganic nitrogen substrate was generated by mixing KNO_3_ and K^15^NO_3_ such that the mixture had an isotopic signature of 0.40 atom % and used directly for the amendments with above mentioned C:N ratio adjustment.

### Elemental and isotopic analysis of plant samples

The shoot and root samples (one mixed sample per pot) that were harvested at the end of the experiment (240h sample for both plant communities) were subjected to carbon and nitrogen elemental and isotope analysis (3 replicates x 3 amendments x 2 plant treatments = 18 samples). Briefly, oven dried shoot and root biomass was ball-milled to a fine powder. Single tin capsules were prepared for total C and N content analysis and analyzed with vario EL CNS analyzer (Elementar Analysensysteme GmBH). For bulk ^13^C and bulk ^15^N analysis finely powdered shoot and root samples were weighed into separate tin capsules. They were analyzed using an elemental analyzer EA1100 (CE, Mainland) coupled via a conflow III interface to a DELTA+ (Finnigan MAT) isotope ratio mass spectrometer (Malik et al., 2015).

### Biomarker extraction, quantification and compound specific isotope analysis

To assess the ^13^C flow into the microbial community, fatty acids from neutral lipids (NL) and phospholipids (PL) were extracted from ingrowth core soil sampled at 0 h and 240 h. All biological replicates of each plant community – soil amendment combination that were collected at the same time point were pooled and homogenized to create a composite sample from which 5g aliquots of soil were used for lipid extraction (1 composite sample x 2 plant community x 3 soil amendments = 6 samples per time point). We applied a modified Bligh and Dyer (1959) lipid extraction protocol using pressurized solvent extraction, as described earlier in Karlowsky et al., 2018a. The concentration of ^13^C incorporated relative to an unlabeled sample (0h) for bacterial PLFA markers (14:0, 15:0, 16:0, 17:0, 18:0, 14:0i, 15:0i, 15:0a, 16:0i, 17:0i, 17:0a, 16:0(10Me), 17:0(10Me), 18:0(10Me), 17:0cy, 19:0cy, 15:1, 16:1ω7, 16:1ω5, 16:1, 17:1, 18:1ω7 and 18:1ω9) and fungal PLFA (18:2ω6,9) was determined using a gas chromatograph coupled with a flame ionization detector (FID) and isotope ratio mass spectrometer (IRMS) respectively (Karlowsky et al., 2018). Additionally, the NLFA biomarker 16:1ω5 (a storage lipid) was used as the marker for AMF.

The amount of ^13^C incorporated was determined from samples collected after the pulse minus the isotope content of unlabeled soil collected before the pulse (Karlowsky et al., 2018). The amounts of saprotrophic fungal markers (PLFA - 18:2ω6,9) were converted into biomass C values by applying conversion factor 11.8 nmol = 1 mg C (Klamer, 2004), for bacterial biomarkers by applying conversion factor 363.3 nmol = 1mg C (Frostegård and Bååth, 1996) and for 16:1ω5 NLFA by applying conversion factor 1.047 nmol = 1μg C (Williams et al., 2017). The amount of ^13^C in the specific biomarker was determined from the C pool size of the biomarker corrected for soil weight and moisture (Karlowsky et al., 2018). The total amount of ^13^C in the bacterial and fungal biomarkers was used as a proxy for microbial biomass that incorporated freshly fixed plant C. Net ^13^C assimilated by soil microbial community integrated across total weight of ingrowth core soil and its parts (saprophytic fungi, AMF and free-living bacteria) for every unit of ^15^N gained by the plant (n=3 per soil amendment) integrated across total above ground biomass was used as an estimate of net plant C allocation for nutrient gain to specific microbial groups.

### DNA and RNA based microbial community profiling

To ascertain the microbial community composition, 1.09 ± 0.01 g (3 replicates x 2 plant treatments x 3 amendments x 1 timepoint (0 hr) = 18 samples) of soil obtained from control samples was used to co-extract total RNA and DNA using the MoBio RNA Powersoil kit. The samples were bead-beaten for one cycle at 5.5 m/s for 30 s with a FastPrep instrument (MP Biomedicals Germany GmbH). DNA was extracted using a MoBio RNA Powersoil DNA elution accessory kit. Traces of DNA that was co-extracted in the final RNA elution were removed using a MoBio DNase Max kit. The purified DNA and RNA concentrations were measured using Qubit 4 fluorometer (Thermo Fisher Scientific) using the broad range assay kits for DNA and RNA. Additionally, the purity of the samples was checked using a nanodrop spectrometer 2000c (Thermo Fisher Scientific) and all samples were found to have A260/280 and A260/230 ratios of 2 suggesting no protein or organic compound contamination. The integrity of the RNA samples was visually verified using 0.8 % agarose gel electrophoresis. Presence of sharp well-defined ribosomal RNA bands and smear in between and above these bands was considered as an indicator of high integrity of the RNA. A replicate was dropped for AM plant community amended with plant root litter and microbial necromass due to poor RNA integrity and subsequent non-amplification of cDNA. 200 ng of total RNA extracted was converted into cDNA using Superscript^™^ VILO kit (Thermo Fisher Scientific) using random hexamers to ensure uniform conversion of all the RNA fragments. The DNA and cDNA samples were further used for amplifying the 16S rRNA gene and ITS regions respectively. Briefly, for the 16S rRNA gene sequencing, amplicon libraries were prepared with V3-V5 primers (Haas et al., 2011) using PCR conditions described by Kozich et al., 2013. One 16S rRNA gene amplicon replicate associated with soil amended with Inorganic-N under NM plant community was dropped due to non-amplification of extracted DNA. ITS2 gene libraries were constructed by amplifying ITS region 2 using the fITS7 (forward) and ITS4 (reverse) primers (Ihrmark et al., 2012) with previously described PCR conditions (Gweon et al., 2015). Both sets of primers also consisted of an appropriate Illumina adapter, an 8-nt index sequence, a 10-nt pad sequence, a 2-nt linker (Kozich et al., 2013). The amplicons were sequenced using Illumina MiSeq, v3 chemistry, 600 cycles.

### Processing of sequencing data

For the bacterial community 16S rRNA gene sequencing data, the forward and reverse reads obtained were merged using PEAR (v 0.9.6) with default parameters (Zhang et al., 2014). The bacterial community composition was analyzed using the QIIME (1.8.0) pipeline (Caporaso et al., 2010). Representative OTU’s were picked using UCLUST using de novo method at 97% similarity level and were assigned taxonomy using RDP classifier method with the SILVA reference data set (release 132 for QIIME) (Edgar, 2010; Wang et al., 2007). Total number of sequencing reads classified within the OTU table for the samples varied between 4000 to 21982 reads. In order to make inter-sample comparisons OTU table obtained was rarefied to 4000 sequences per sample using QIIME command single_rarefraction.py (Caporaso et al., 2010).

Since fungal spores are ubiquitous, the ITS2 regions that were transcribed into RNA were used to sample transcriptionally active fungi (Anderson and Parkin, 2007). The amplified ITS region from total cDNA samples were used for sequencing. The amplified ITS sequencing data was analyzed using the PIPITS pipeline using the default parameters (Gweon et al., 2015). Briefly, the paired end sequences were merged using VSEARCH after which the ITS2 region was extracted using ITSx after which they are matched to a trained UNITE database at 97% similarity using RDP classifier to obtain an OTU table (Bengtsson-Palme et al., 2013; Rognes et al., 2016). The OTUs detected were then assigned to fungal guilds using FUNguild assignment app (http://www.stbates.org/guilds/app.php) (Nguyen et al., 2016). The proportion of ITS community assigned to AMF was also used as evidence of AMF regrowth/presence in the ingrowth core.

The bacterial and fungal amplicon sequencing data and the associated metadata is publicly available through the Sequence Read Archive project number: SRP132171. A detailed table of associated metadata and download links is also provided in the supplementary information (Table S12 and S13).

### Statistical data analysis

One-way ANOVA was used to determine statistical differences amongst rhizosphere types (AM vs NM) for root:shoot ratio, ^15^N assimilated by plant shoot and net microbial ^13^C assimilation into total microbial biomass. Two-way ANOVA was applied to identify statistically significant differences due to rhizosphere and litter types for plant carbon allocation per unit ^15^N gain. All two-way ANOVA models that showed significant interactions of rhizosphere-litter types were subject to Tukey’s post-hoc test to identify differences amongst litter types nested within rhizosphere types. Significant differences were only considered if the assumption for homogeneity of variance was met using the Flinger-Killing test (Table S2). Subsampled OTU tables were binned to different taxonomic groups at the phylum and class levels (bacteria) whereas fungal OTUs were binned into guilds to compare effect of AM vs NM plant communities and substrate amendments on microbial (bacterial and fungal) community composition. The effects of substrate and plant community on microbial groups was determined using a two-way ANOVA of their relative abundance. Differences between microbial (bacterial and fungal) community composition affected by the experimental treatment were determined using non-parametric Permutational Multivariate Analysis of Variance PERMANOVA (“adonis” function in the vegan package 2.5-2) (Oksanen et al., 2018). The assumption of homogeneity of dispersions (variance) was tested using the betadisper function (vegan package 2.5-2) and both the fungal and bacterial community composition was found to meet this assumption (Table S9). The dissimilarity based upon microbial community composition due to litter and rhizosphere treatments were visualized as an NMDS ordination of Bray-Curtis dissimilarity coefficients was plotted to visualize the dissimilarities between the microbial communities under the influence of plant communities and substrate added to the soil. All analysis was performed using R software (v 3.4.3) with visualization being done with ggplot2 package (Wickham, 2011).

## Results

### Carbon allocation and Nitrogen return

The AM plant community had a lower shoot biomass (4.62 ± 0.14 g, n = 9 aggregated across substrate amendments) compared to NM the plant community (7.42 ± 0.44 g, n = 9 aggregated across substrate amendments) but both had similar root biomass (F_*1,16*_ = 0.17; p = 0.685), which resulted in a higher root to shoot ratio for the AM plant community. The AM plant community also allocated more carbon into their roots, which was reflected in the root to shoot ratio of ^13^C incorporated into the AM plant community compared to the NM plant community (F_*1,16*_ = 20.91, p < 0.001; Figure 2A). Unexpectedly, the net ^13^C assimilation of the rhizosphere microbial communities from the AM plant community was significantly lower than that of the NM plant community (F_*1,16*_ = 21.75, p < 0.001, Figure 2B).

**Figure 2:**
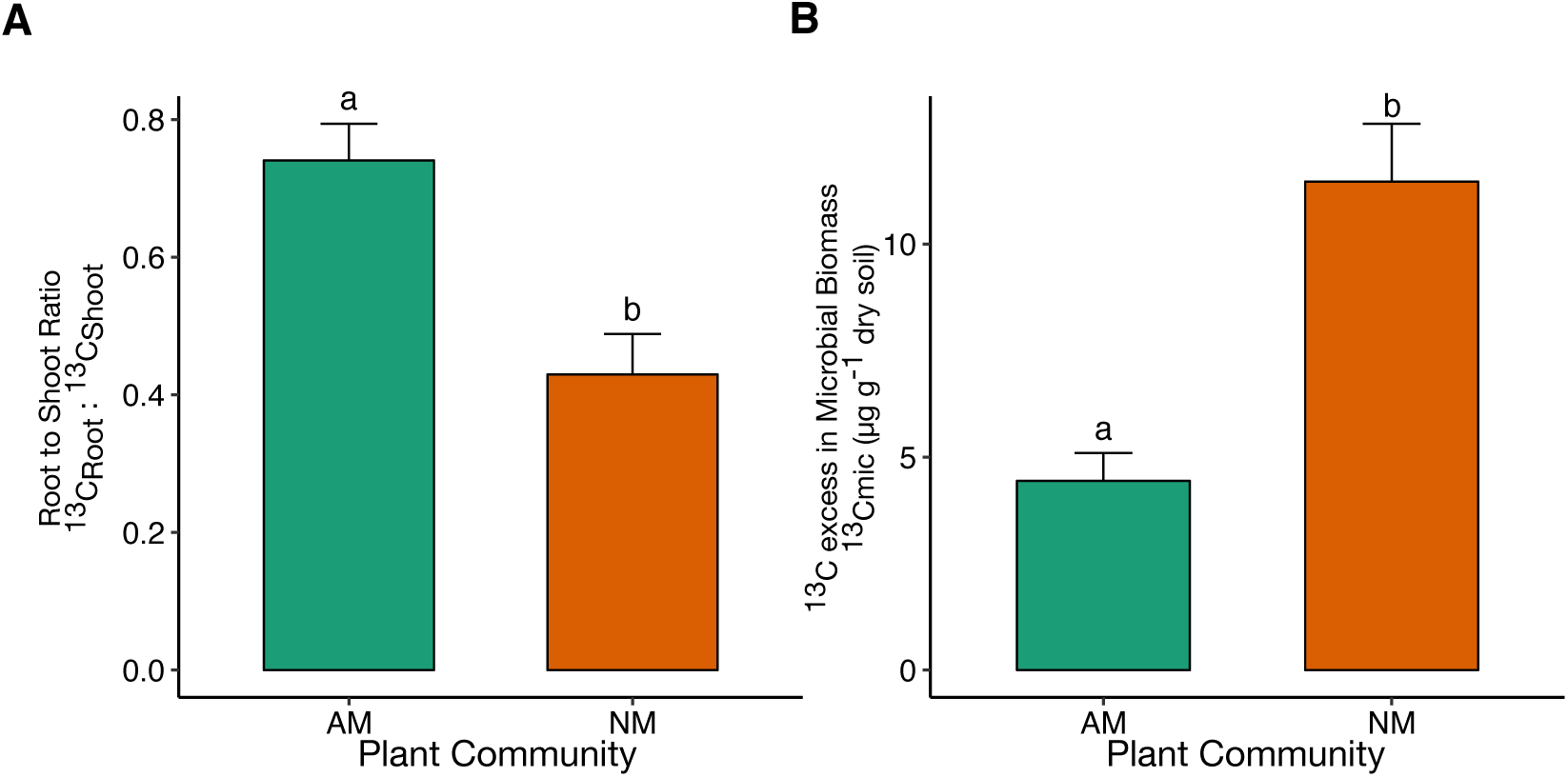
Photosynthetically fixed carbon assimilated by plants and associated microbial community. **A:** Comparison of root to shoot ^13^C content between AM (n = 9) and NM (n = 9) the plant community types across all soil amendments. **B:** Mean ^13^C (ng) excess in soil microbial biomass of each plant community types. Height of the bars represents mean and error bars represent standard error of mean. Different letters on bars designate significant differences based on one way analysis of variance (ANOVA) comparing plant community types. Colors represent plant community types where Dark Green=AM plant community and Orange= NM plant community.

The AM plant community gained 93.1 ± 4.7 μg ^15^N from litter decomposition, which was not significantly different from the NM plant community that gained 82.3 ± 10.7 μg ^15^N ((F_1,16_ = 0.40, p = 0.541, Figure 3A). A comparison across substrate amendments nested within either plant community showed no significant differences in terms of ^15^N uptake (AM: F_*2,6*_ = 0.57, p = 0.594; NM: F_*2,6*_ = 0.797, p = 0.493). However, the AM plant community invested 66% less ^13^C into the microbial community per ^15^N gain than the NM plant community (F_*1,16*_ = 15.20, p = 0.001; Figure 3B). Moreover, the AM plant community allocated significantly more ^13^C per ^15^N gained to AMF biomass (F_*1,16*_ = 30.53, p < 0.001; Figure 4A) whereas NM plant communities allocated significantly more ^13^C per ^15^N gained into saprophytic fungi (F_*1,16*_ = 15.47, p = 0.001; Figure 4B) and bacteria (F_*1,16*_ =4.78, p=0.044). Unexpectedly the amount of ^13^C allocated per ^15^N gained depended only in the AM plant community on the type of decomposed substrate (F_2,15_ = 7.13, p = 0.009, Figure 4).

**Figure 3:**
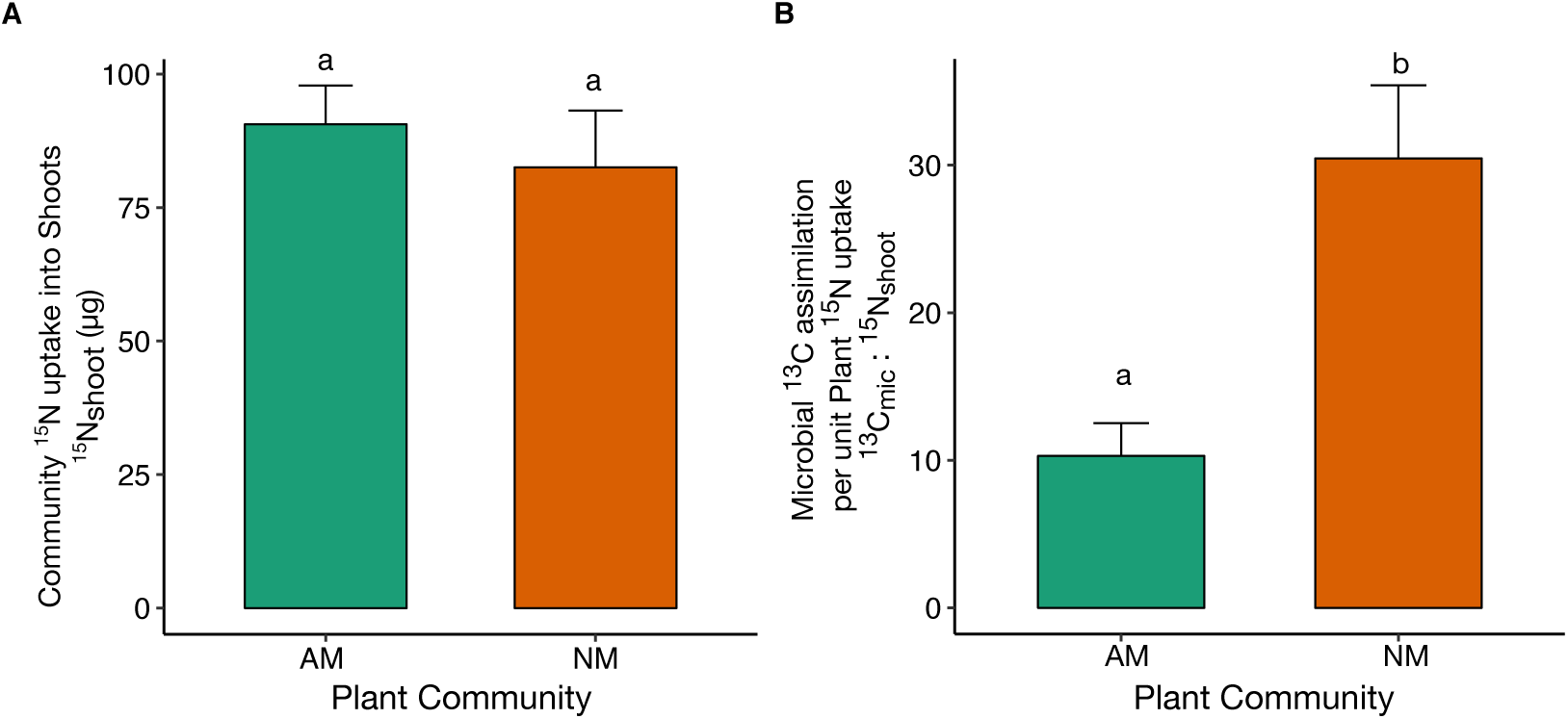
Carbon investment into microbial community and return of decomposition released nitrogen. **A:** ^15^N content (µg) of shoot biomass compared between AM (n = 9) and NM (n = 9) for plant communities. **B:** Comparison of carbon investment by plant communities into total microbial biomass as determined from ^13^C excess in total microbial biomass (PLFA) scaled up to ingrowth core soil weight (200 g) per μg for respective plant communities. Colors represent plant community types (Dark Green = AM and Orange = NM plant community). In both cases, soil amendments were aggregated for each plant community. Horizontal bars represent means for each group and error bars represent standard error of mean across all soil amendment for the specific plant community. Groups sharing the same letters are not significantly different based on one-way analysis of variance (ANOVA) comparing only plant communities.

**Figure 4:**
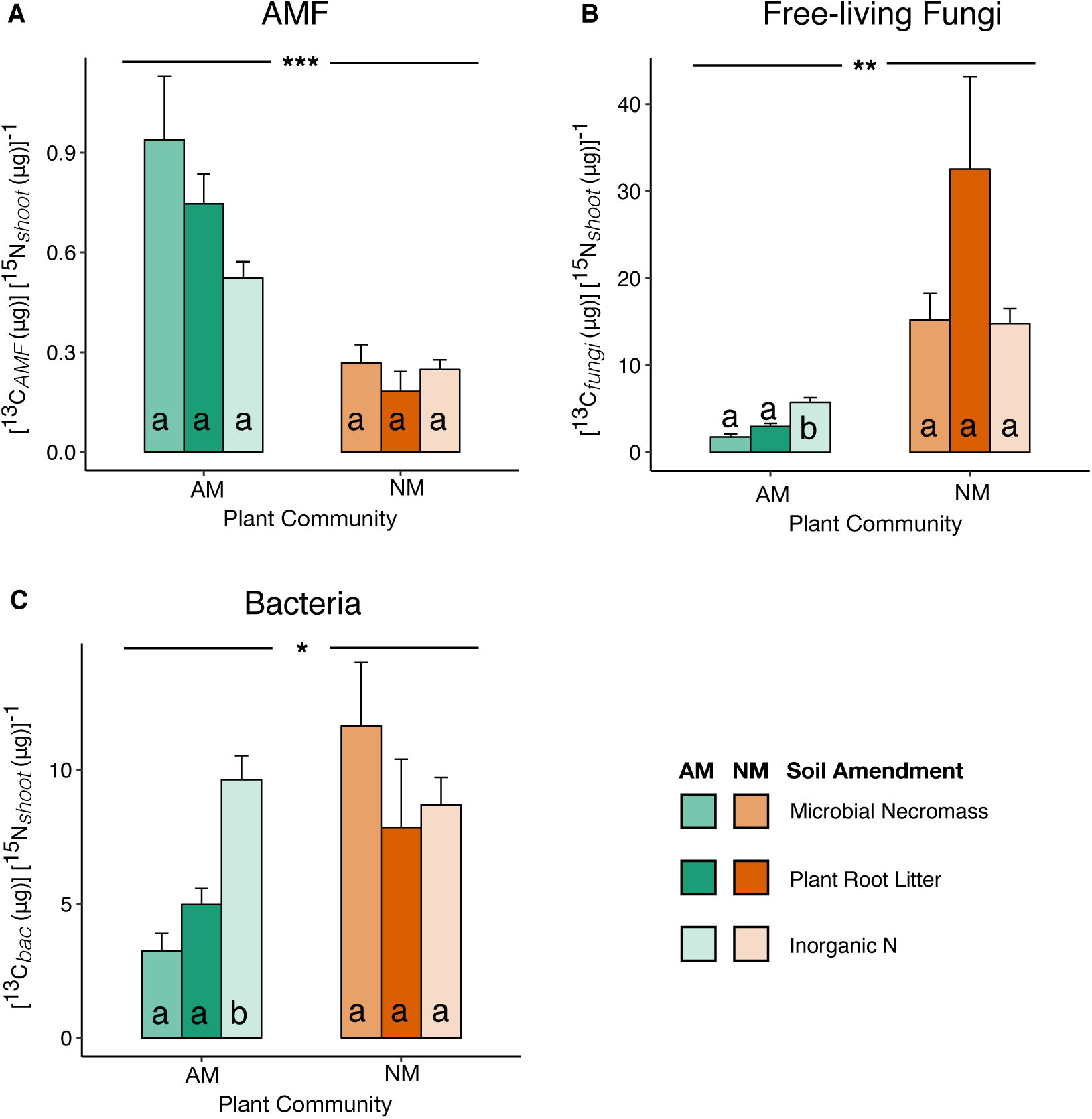
^13^C investment for ^15^N gain by plants partitioned based on microbial groups. Mean ^13^C incorporated into total microbial group specific lipids (A – AMF (NLFA) ; B – Free-living fungi (PLFA) and C – Bacteria (PLFA)) per unit ^15^N uptake by the plant community. Standard error calculated based on replicated plant community ^15^N uptake measurements (n = 3 per soil amendment nested under plant community type). Means, standard error and significant differences between plant communities (asterisk) and soil amendments (LSD letter-group based on Tukey’s post-hoc tests) are shown. Significance is indicated at p < 0.05 (*), p < 0.01 (**) and p <0.001 (***).

Specifically, in the AM rhizosphere, colonization of inorganic N amended soil resulted in significantly more plant-derived ^13^C to be allocated to net bacteria and saprotrophic fungal biomass compared to microbial necromass and plant litter amended soil (Figure 4B, C). Such litter-based differences were missing in the NM rhizosphere.

### Microbial community composition

The microbial communities fostered by presence or absence of AMF associations were similar in terms of richness and evenness (Table S3 and S4). However, individual bacterial groups and fungal guilds were responsive to either presence or absence of AMF association (AM and NM) or to specific substrate amendments (plant root litter, microbial necromass and inorganic-N). Presence of AMF association significantly increased relative abundance of bacterial groups belonging to *Actinobacteria* and *Proteobacteria* (specifically *Betaproteobacteria)* (Table S5) whereas absence of AMF associations significantly increased the abundance of the *Acidobacteria* group (Table S5). In agreement with results from PLFA analysis, absence of AMF association also significantly increased reliance of the plant community on active fungal saprotrophs and endophytes (Table S6), both of which are guilds that can lead a free-living lifestyles (Rodriguez et al., 2009). In contrast and as expected, the presence of AMF associations resulted in significantly higher relative abundance of symbiotrophic AMF guilds (Table S6) and reduced abundance of free-living fungal guilds (Saprotrophs and Endophytes). As the fungal ITS community was determined using cDNA-based amplicon sequencing, dominance of AMF along with copious root ingrowth (Figure S2) was also used as confirmatory evidence of AMF colonization of ingrowth core soil.

Substrate amendments on the other hand elicited specific responses in different bacterial groups depending on presence or absence of AMF associations. Bacterial groups belonging to *Gemmamonadetes, Nitrospira* and *Deltaproteobacteria* were observed to be significantly affected by substrate amendments (Table S7). In all cases, the differences could be consistently explained by substrate source, with higher abundances of these groups observed in ingrowth cores amended with plant root litter relative to other amendments. The fungal OTUs that were not exclusively assigned to major guilds (Saprotrophs, AMF, Ectomycorrhizal and Endophytes) were observed to show a significant response to substrate amendments (Table S8).

Fungal community composition varied on expected lines with presence or absence of AMF association explaining dissimilarity amongst microbial communities (Figure 5A). Ordinations of fungal communities also revealed that effects of substrate amendments were nested within plant community treatment (Figure 5A). Ingrowth core fungal community composition in presence of AMF association depended mainly on type of nitrogen source with clear separation of communities between organic and inorganic nitrogen sources (Figure 5A). On the other hand, ingrowth core microbial communities in absence of AMF association were separated by substrate origin with a clear separation between plant root litter and remaining amendments (Figure 5A). Bacterial community compositions were largely dependent on substrate source (Figure 5B). Within these substrate amendments AMF association resulted in sub clusters for plant root litter and inorganic N amendments only (Figure 5B).

**Figure 5:**
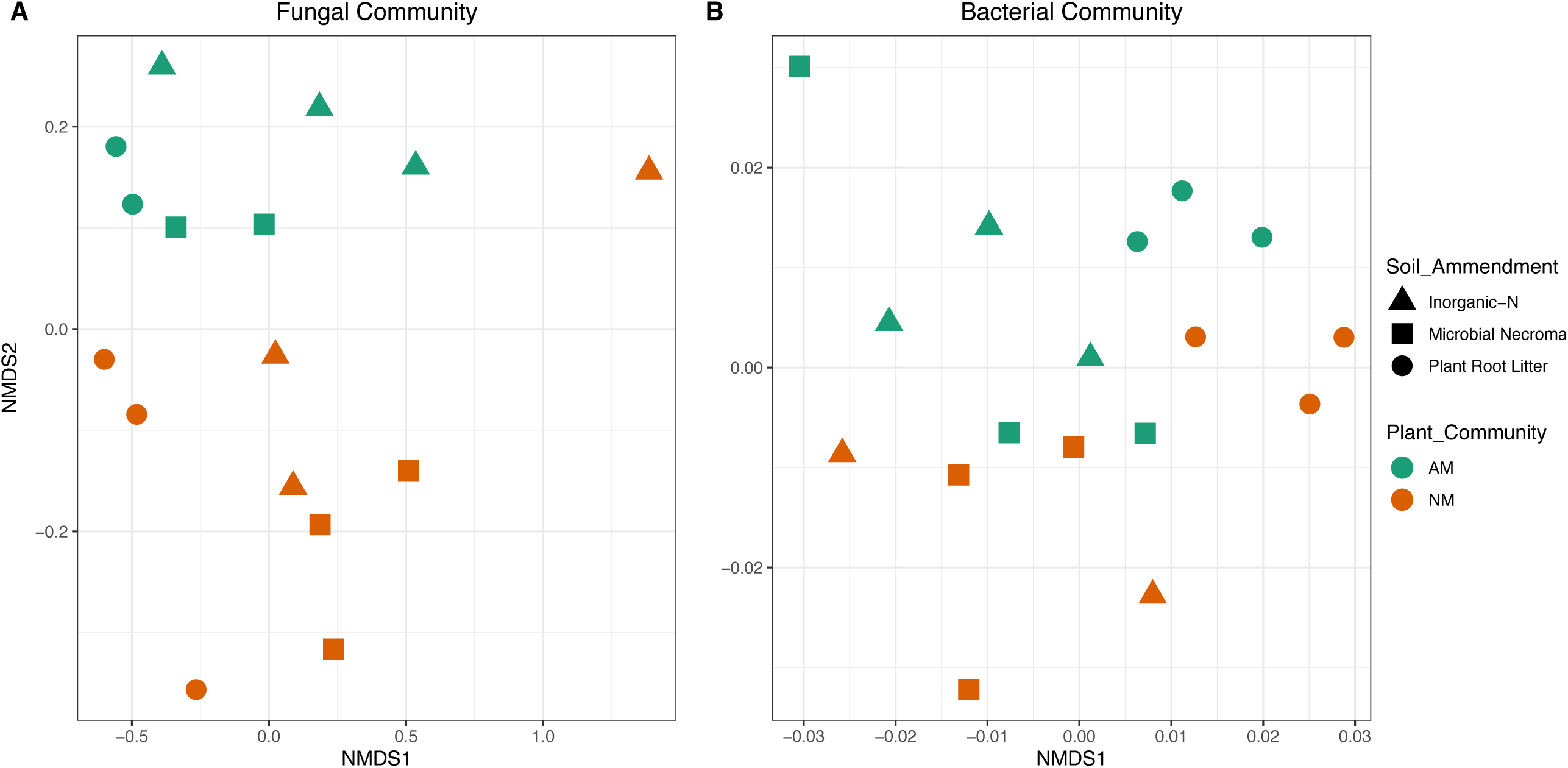
Variation in microbial bacterial and fungal community composition. Non-metric multidimensional scaling (NMDS) plots illustrating Bray-Curtis distances between microbial communities (A: Fungal Community and B: Bacterial Community) in individual samples. Colors represent plant community type (Dark Green = AM and Orange = NM) whereas symbols represents soil amendments. Stress values for fungal (B) and bacterial (A) plots are 0.05 and 0.18 respectively.

Permutational Multivariate Analysis of variance (PERMANOVA) revealed that both fungal (Table S10) and bacterial community (Table S11) compositions were significantly affected by plant community and substrate amendments. In terms of variance explained by presence or absence of AMF association, fungal community composition was affected more than bacterial community. In both bacterial and fungal community substrate amendments explained a greater proportion of the variance in community composition.

## Discussion

In this study we investigated how plant communities, differing in their association with AMF, affected the carbon allocation to the microbial communities and the return of mineralized nitrogen from different substrates. By combining a dual stable isotope labelling with a sequencing analyses and compound specific isotope analysis of the soil microbial community we found that AMF associated (AM) plant communities did not gain significantly more nitrogen from the different substrates than non-mycorrhizal (NM) plant communities. However, AM plant communities invested less carbon into the soil microbial communities than NM communities for similar nutrient uptake (Figure 6), which was confirmed by total microbial biomass (chloroform fumigation) and soil respiration (Figure S4 and S5). Even though AM plant communities foster a carbon demanding fungal network, they benefit strongly from the AMF’s targeted access to the decomposition sites where plant-derived carbon is used to stimulate selected microbial groups for substrate decomposition. This important role of the hyphae is underscored by the fact that a comparison of compartments with only hyphae access (30µm mesh) and root and hyphae access (100 µm mesh) did not differ in their performance (Table S1). These findings are also in line with previous studies showing the importance of the AMF hyphal network on the associated soil bacterial community (Drigo et al., 2010; Vestergård et al., 2008). The close interaction of AMF and bacteria potentially increases decomposition efficiency and makes in the long run more nutrients available to the plant communities. NM plant communities likely miss the targeted growth of AMF hyphae and rely on free-living saprophytic fungi and bacteria for decomposition, therefore increasing the overall carbon per nitrogen cost.

**Figure 6.**
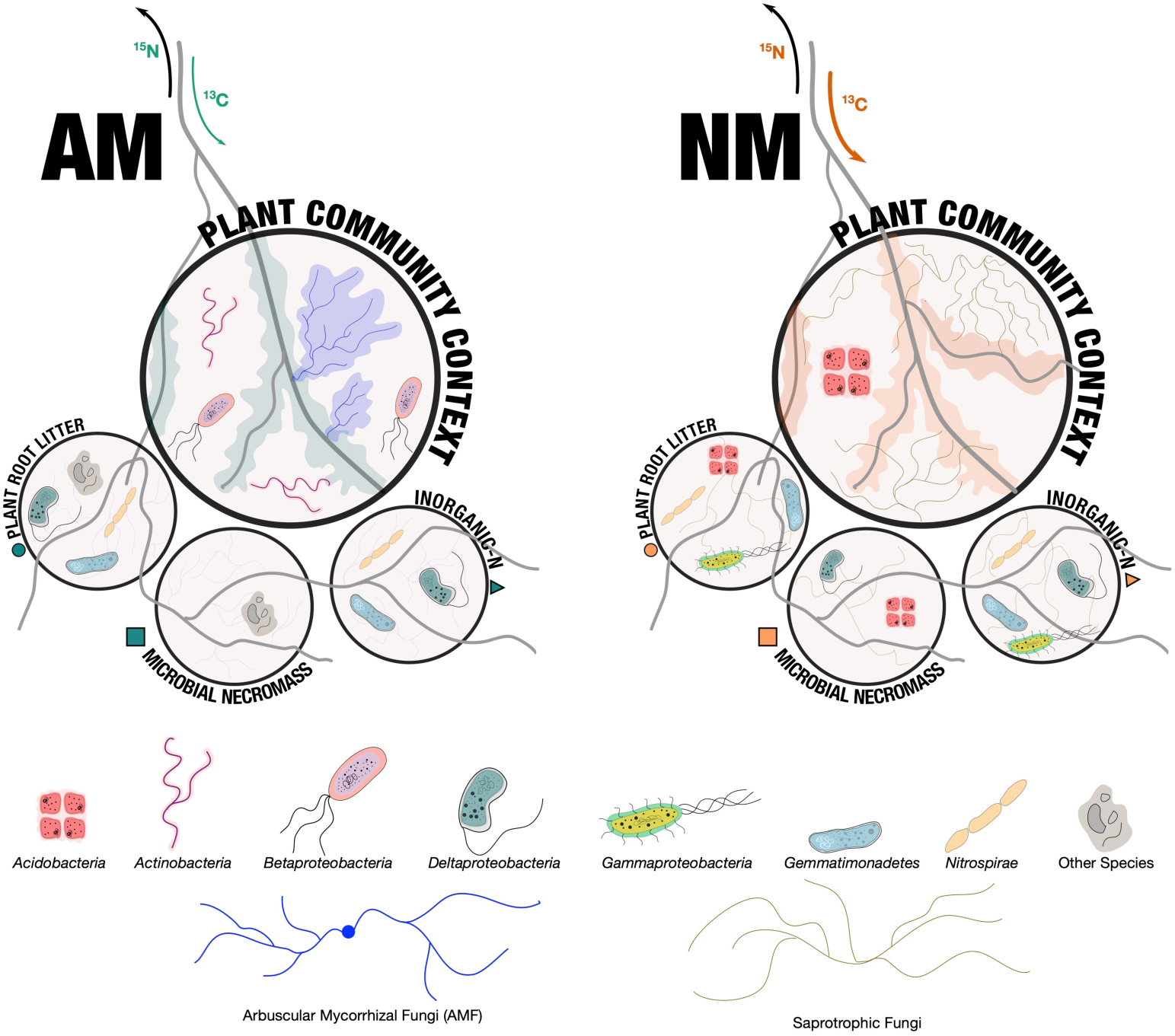
Conceptual model showing the effect of AMF association on decomposer and plant responsive microbial groups that shape carbon and nutrient flux when decomposing stoichiometrically identical substrates. In non-mycorrhizal plants (NM) the role of AMF is taken over by fungal saprotrophs. Distinct bacterial communities are assembled for each substrate depending on substrate origin and plant-AMF association context. Arrows represent the C and N flow differences arising from presence or absence of AMF, explaining that for an equivalent N uptake AMF associated plants invested lower amounts of photoassimilated carbon into decomposer community.

Furthermore, our results also showed that the net assimilation of plant carbon by the rhizosphere microbial community depended on the biochemical characteristics of the substrates available for decomposition. This strongly suggests that substrate (bio)chemistry is as important as nutrient content of the substrate in the decomposition process and can modulate the extent of plant-derived carbon required for decomposition (Jones et al., 2004). AMF-associated rhizosphere microbial communities generally assimilated less plant carbon than the microbial communities lacking this association, which suggests that AMF allocated less labile carbon to the free-living soil microbial community in order to take up mineralized N from decomposition processes in soil. The different net assimilation of the free-living microbial community from AM and NM plant communities also suggests that the decomposition of different substrates strongly depended on the association with AMF. This confirms that AMF indeed act as target-oriented root extensions that more effectively feed plant-derived carbon to the free-living microbial community. Moreover, AMF through their response to substrate biochemistry and nutrient availability likely control the amount of exudates released for supporting decomposition related processes. In turn, the significant response of specific microbial groups to either the litter type and rhizosphere type as well as the shift in the microbial community composition suggests that in addition to the substrate nutrient content (Stempfhuber et al., 2016) the interplay of substrate source with plant carbon input also shapes microbial communities in the rhizosphere. Hence the composition of the microbial decomposer community is adjusted to plant exudation and type of the substrate being decomposed (Figure 6), making AMF associations with plants an important part of the decomposition process (McLaren and Turkington, 2011). Expectedly, the fungal community primarily differed between AM or NM plant communities, emphasizing the fungal role in decomposition and nutrient gathering (Bukovská et al., 2018; Meier et al., 2015). However, in the presence of AMF, we observed lower plant carbon assimilation by the free living microbial community (Mueller et al., 2014), indicating dominance of bacterial community in the associated decomposition processes. This suggests that primary decomposers in the AM rhizosphere were bacteria that were fed and adapted to substrate types rapidly by the AMF (Herman et al., 2012; Nuccio et al., 2013). Most interestingly our results indicate for the first time that AM plant communities strongly benefit from the association with AMF during nutrient uptake from microbial necromass. AMF assimilated more ^13^C from plants but invested less carbon into the bacterial community for decomposition. This might indicate that AMF, which have no capability to decompose lignocellulose structures of plant cell walls themselves (Tisserant et al., 2013), might be able to decompose peptidoglycan structures of microbial cell walls. This idea would be supported by the dramatic up-regulation of genes encoding for enzymes containing a peptidoglycan binding LysM domain and glycan hydrolases like lysozyme, glucosamidase and alpha amylase that would be necessary for microbial cell wall decomposition (Tisserant et al., 2013; Gude et al., 2012). If we consider that large parts of soil organic matter are microbial necromass (Gleixner, 2013) the association of plants with AMF gain a new overall importance for ecosystem functioning. However, additional experiments are needed to confirm this new role of plant-AMF associations for microbial cell wall decomposition.

## Conclusion

Our results suggest that a three-way interaction between the plant community, the microbial community and the type of decomposed substrate is of fundamental importance for ecosystem functioning. Specifically, nutrient recycling in the plant-microorganism-soil continuum is shaped by these interactions leading to a plant-driven and context-dependent stimulation of decomposers whose decomposition capabilities are likely versatile as stoichiometrically identical litter differing in its origin are decomposed by different parts of the microbial community. Consequently, organic matter decomposition and nutrient recycling can only be fully understood if the whole continuum is investigated. Moreover, our results demonstrate for the first time that plant nutrient gain via AMF association is less carbon intensive and likely more targeted than passive exudation of plant communities lacking an AMF association. This might explain why a large part of the plant community rely on AMF mediated nutrient uptake strategy.

## Acknowledgements

SC received funding from long term DAAD scholarship to carry out the research. ML is funded by the German Research Foundation (DFG; FOR 456, FOR 1451 – “The Jena Experiment”) and by the “Zwillenberg-Tietz Stiftung”. We acknowledge help from Agnes Fastnacht with greenhouse resources and monitoring of the experiment. Special thanks to Karl Kübler for construction and deployment of the pulse labelling setup in the greenhouse. We acknowledge Heike Geilmann and Steffen Ruehlow for help with stable isotope measurements, and Maria Foerster for helping with fatty acid analysis We also thank Prof. (Dr.) Erika Kothe, Ruchira Mukherji, Elisa Catao and Huei Ying Gan for helpful comments and discussions.

## Author contributions

SC, GG and AM designed the study. SC and JH performed the experiment. SC performed the extractions and stable isotope analyses. TG and RG completed the downstream processing of nucleic acids, sequencing and supervised bioinformatics analysis. SC and ML completed the statistical analysis. SC, ML and GG conceptualized the work and wrote the manuscript and all authors contributed to revisions.

## References

Anderson, I. C., and Parkin, P. I. (2007). Detection of active soil fungi by RT-PCR amplification of precursor rRNA molecules. J. Microbiol. Methods 68, 248–253. doi:10.1016/j.mimet.2006.08.005.

Averill, C., Bhatnagar, J. M., Dietze, M. C., Pearse, W. D., and Kivlin, S. N. (2019). Global imprint of mycorrhizal fungi on whole-plant nutrient economics. Proc. Natl. Acad. Sci. U. S. A. 116, 23163–23168. doi:10.1073/pnas.1906655116.

Bengtsson-Palme, J., Ryberg, M., Hartmann, M., Branco, S., Wang, Z., Godhe, A., et al. (2013). Improved software detection and extraction of ITS1 and ITS2 from ribosomal ITS sequences of fungi and other eukaryotes for analysis of environmental sequencing data. Methods Ecol. Evol. 4, 914–919. doi:10.1111/2041-210X.12073.

Bligh, E. G., and Dyer, W. J. (1959). A rapid method of total lipid extraction and purification. Can. J. Biochem. Physiol. 37, 911–917. doi:10.1139/y59-099.

Bonfante, P., and Genre, A. (2010). Mechanisms underlying beneficial plant-fungus interactions in mycorrhizal symbiosis. Nat. Commun. 1, 48. doi:10.1038/ncomms1046.

Bowles, T. M., Jackson, L. E., and Cavagnaro, T. R. (2018). Mycorrhizal fungi enhance plant nutrient acquisition and modulate nitrogen loss with variable water regimes. Glob. Chang. Biol. 24, e171–e182. doi:10.1111/gcb.13884.

Brundrett, M.. (2015a). Arbuscular Mycorrhizas. In: Mycorrhizal Associations: The Web Resource. Version 2.0. 24.04.2015.‹mycorrhizas.info›. Available at: http://mycorrhizas.info/vam.html.

Brundrett, M.. (2015b). Mycorrhizal Associations: The Web Resource. Version 2.0. 24.04.2015. ‹mycorrhizas.info›. Available at: http://mycorrhizas.info/nmplants.html.

Bukovská, P., Bonkowski, M., Konvalinková, T., Beskid, O., Hujslová, M., Püschel, D., et al. (2018). Utilization of organic nitrogen by arbuscular mycorrhizal fungi—is there a specific role for protists and ammonia oxidizers? Mycorrhiza 28, 269–283. doi:10.1007/s00572-018-0825-0.

Bunn, R. A., Simpson, D. T., Bullington, L. S., Lekberg, Y., and Janos, D. P. (2019). Revisiting the ‘direct mineral cycling’ hypothesis: arbuscular mycorrhizal fungi colonize leaf litter, but why? ISME J., 1. doi:10.1038/s41396-019-0403-2.

Caporaso, J. G., Kuczynski, J., Stombaugh, J., Bittinger, K., Bushman, F. D., Costello, E. K., et al. (2010). QIIME allows analysis of high-throughput community sequencing data. Nat. Methods 7, 335–336. doi:10.1038/nmeth.f.303.

Craine, J. M., Morrow, C., and Fierer, N. (2007). Microbial nitrogen limitation increases decomposition. Ecology 88, 2105–2113. doi:10.1890/06-1847.1.

Davison, J., Moora, M., Opik, M., Adholeya, A., Ainsaar, L., Ba, A., et al. (2015). Global assessment of arbuscular mycorrhizal fungus diversity reveals very low endemism. Science 349, 970–973. doi:10.1126/science.aab1161.

de Graaff, M.-A., Classen, A. T., Castro, H. F., and Schadt, C. W. (2010). Labile soil carbon inputs mediate the soil microbial community composition and plant residue decomposition rates. New Phytol. 188, 1055–1064. doi:10.1111/j.1469-8137.2010.03427.x.

Dijkstra, F. A., Carrillo, Y., Pendall, E., and Morgan, J. A. (2013). Rhizosphere priming: a nutrient perspective. Front. Microbiol. 4, 216. doi:10.3389/fmicb.2013.00216.

Drigo, B., Pijl, A. S., Duyts, H., Kielak, A. M., Gamper, H. A., Houtekamer, M. J., et al. (2010). Shifting carbon flow from roots into associated microbial communities in response to elevated atmospheric CO2. Proc. Natl. Acad. Sci. U. S. A. 107, 10938–42. doi:10.1073/pnas.0912421107.

Edgar, R. C. (2010). Search and clustering orders of magnitude faster than BLAST. Bioinformatics 26, 2460–2461. doi:10.1093/bioinformatics/btq461.

Frostegård, A., and Bååth, E. (1996). The use of phospholipid fatty acid analysis to estimate bacterial and fungal biomass in soil. Biol. Fertil. Soils 22, 59–65. doi:10.1007/s003740050076.

Gleixner, G. (2013). Soil organic matter dynamics: a biological perspective derived from the use of compound-specific isotopes studies. Ecol. Res. 28, 683–695. doi:10.1007/s11284-012-1022-9.

Güsewell, S., and Gessner, M. O. (2009). N:P ratios influence litter decomposition and colonization by fungi and bacteria in microcosms. Funct. Ecol. 23, 211–219. doi:10.1111/j.1365-2435.2008.01478.x.

Gweon, H. S., Oliver, A., Taylor, J., Booth, T., Gibbs, M., Read, D. S., et al. (2015). PIPITS: An automated pipeline for analyses of fungal internal transcribed spacer sequences from the Illumina sequencing platform. Methods Ecol. Evol. 6, 973–980. doi:10.1111/2041-210X.12399.

Haas, B. J., Gevers, D., Earl, A. M., Feldgarden, M., Ward, D. V, Giannoukos, G., et al. (2011). Chimeric 16S rRNA sequence formation and detection in Sanger and 454-pyrosequenced PCR amplicons. Genome Res. 21, 494–504. doi:10.1101/gr.112730.110.

Harley, J. L., and Harley, E. L. (1987). A Checklist of Mycorrhiza in the British Flora. New Phytol. 105, 1–102. doi:10.1111/j.1469-8137.1987.tb00674.x.

Herman, D. J., Firestone, M. K., Nuccio, E., and Hodge, A. (2012). Interactions between an arbuscular mycorrhizal fungus and a soil microbial community mediating litter decomposition. FEMS Microbiol. Ecol. 80, 236–247. doi:10.1111/j.1574-6941.2011.01292.x.

Hestrin, R., Hammer, E. C., Mueller, C. W., and Lehmann, J. (2019). Synergies between mycorrhizal fungi and soil microbial communities increase plant nitrogen acquisition. Commun. Biol. 2, 233. doi:10.1038/s42003-019-0481-8.

Husson, O. (2013). Redox potential (Eh) and pH as drivers of soil/plant/microorganism systems: A transdisciplinary overview pointing to integrative opportunities for agronomy. Plant Soil 362, 389–417. doi:10.1007/s11104-012-1429-7.

Hütsch, B. W., Augustin, J., and Merbach, W. (2002). Plant rhizodeposition — an important source for carbon turnover in soils. J. Plant Nutr. Soil Sci. 165, 397. doi:10.1002/1522-2624(200208)165:4<397::AID-JPLN397>3.0.CO;2-C.

Ihrmark, K., Bödeker, I. T. M., Cruz-Martinez, K., Friberg, H., Kubartova, A., Schenck, J., et al. (2012). New primers to amplify the fungal ITS2 region - evaluation by 454-sequencing of artificial and natural communities. FEMS Microbiol. Ecol. 82, 666–677. doi:10.1111/j.1574-6941.2012.01437.x.

Jacoby, R., Peukert, M., Succurro, A., Koprivova, A., and Kopriva, S. (2017). The Role of Soil Microorganisms in Plant Mineral Nutrition-Current Knowledge and Future Directions. Front. Plant Sci. 8, 1617. doi:10.3389/fpls.2017.01617.

Jilling, A., Keiluweit, M., Contosta, A. R., Frey, S., Schimel, J., Schnecker, J., et al. (2018). Minerals in the rhizosphere: overlooked mediators of soil nitrogen availability to plants and microbes. Biogeochemistry 139, 103–122. doi:10.1007/s10533-018-0459-5.

Jones, D. L., Hodge, A., and Kuzyakov, Y. (2004). Plant and mycorrhizal regulation of rhizodeposition. New Phytol. 163, 459–480. doi:10.1111/j.1469-8137.2004.01130.x.

Kaiser, C., Kilburn, M. R., Clode, P. L., Fuchslueger, L., Koranda, M., Cliff, J. B., et al. (2015). Exploring the transfer of recent plant photosynthates to soil microbes: Mycorrhizal pathway vs direct root exudation. New Phytol. 205. doi:10.1111/nph.13138.

Kallenbach, C. M., Frey, S. D., and Grandy, A. S. (2016). Direct evidence for microbial-derived soil organic matter formation and its ecophysiological controls. Nat. Commun. 7, 13630. doi:10.1038/ncomms13630.

Karlowsky, S., Augusti, A., Ingrisch, J., Hasibeder, R., Lange, M., Lavorel, S., et al. (2018). Land use in mountain grasslands alters drought response and recovery of carbon allocation and plant-microbial interactions. J. Ecol. 106, 1230–1243. doi:10.1111/1365-2745.12910.

Keiluweit, M., Bougoure, J. J., Nico, P. S., Pett-Ridge, J., Weber, P. K., and Kleber, M. (2015). Mineral protection of soil carbon counteracted by root exudates. Nat. Clim. Chang. 5, 588–595. doi:10.1038/nclimate2580.

Klamer, M. (2004). Estimation of conversion factors for fungal biomass determination in compost using ergosterol and PLFA 18:2ω6,9. Soil Biol. Biochem. 36, 57–65. doi:10.1016/j.soilbio.2003.08.019.

Kozich, J. J., Westcott, S. L., Baxter, N. T., Highlander, S. K., and Schloss, P. D. (2013). Development of a dual-index sequencing strategy and curation pipeline for analyzing amplicon sequence data on the miseq illumina sequencing platform. Appl. Environ. Microbiol. 79, 5112–5120. doi:10.1128/AEM.01043-13.

Kuzyakov, Y., and Xu, X. (2013). Competition between roots and microorganisms for nitrogen: mechanisms and ecological relevance. New Phytol. 198, 656–69. doi:10.1111/nph.12235.

Levin, P. A., and Angert, E. R. (2015). Small but mighty: Cell size and bacteria. Cold Spring Harb. Perspect. Biol. 7, 1–11. doi:10.1101/cshperspect.a019216.

Liang, C., Schimel, J. P., and Jastrow, J. D. (2017). The importance of anabolism in microbial control over soil carbon storage. Nat. Microbiol. 2, 17105. doi:10.1038/nmicrobiol.2017.105.

Malik, A. A., Dannert, H., Griffiths, R. I., Thomson, B. C., and Gleixner, G. (2015). Rhizosphere bacterial carbon turnover is higher in nucleic acids than membrane lipids: Implications for understanding soil carbon cycling. Front. Microbiol. 6, 268. doi:10.3389/fmicb.2015.00268.

Malik, A. A., Puissant, J., Buckeridge, K. M., Goodall, T., Jehmlich, N., Chowdhury, S., et al. (2018). Land use driven change in soil pH affects microbial carbon cycling processes. Nat. Commun. 9, 1–10. doi:10.1038/s41467-018-05980-1.

McLaren, J. R., and Turkington, R. (2011). Plant identity influences decomposition through more than one mechanism. PLoS One 6, e23702. doi:10.1371/journal.pone.0023702.

Meier, I. C., Pritchard, S. G., Brzostek, E. R., McCormack, M. L., and Phillips, R. P. (2015). The rhizosphere and hyphosphere differ in their impacts on carbon and nitrogen cycling in forests exposed to elevated CO_2_. New Phytol. 205, 1164–74. doi:10.1111/nph.13122.

Mueller, R. C., Balasch, M. M., and Kuske, C. R. (2014). Contrasting soil fungal community responses to experimental nitrogen addition using the large subunit rRNA taxonomic marker and cellobiohydrolase I functional marker. Mol. Ecol. 23, 4406–17. doi:10.1111/mec.12858.

Nguyen, N. H., Song, Z., Bates, S. T., Branco, S., Tedersoo, L., Menke, J., et al. (2016). FUNGuild: An open annotation tool for parsing fungal community datasets by ecological guild. Fungal Ecol. 20, 241–248. doi:10.1016/J.FUNECO.2015.06.006.

Nuccio, E. E., Hodge, A., Pett-Ridge, J., Herman, D. J., Weber, P. K., and Firestone, M. K. (2013). An arbuscular mycorrhizal fungus significantly modifies the soil bacterial community and nitrogen cycling during litter decomposition. Environ. Microbiol. 15, 1870–1881. doi:10.1111/1462-2920.12081.

Nuccio, E. E., Starr, E., Karaoz, U., Brodie, E. L., Zhou, J., Tringe, S. G., et al. (2020). Niche differentiation is spatially and temporally regulated in the rhizosphere. ISME J. 14, 999–1014. doi:10.1038/s41396-019-0582-x.

Oksanen, J., Blanchet, F. G., Friendly, M., Kindt, R., Legendre, P., McGlinn, D., et al. (2018). vegan: Community Ecology Package. Available at: https://cran.r-project.org/package=vegan.

Rodriguez, R. J., White Jr, J. F., Arnold, A. E., and Redman, R. S. (2009). Fungal endophytes: diversity and functional roles. New Phytol. 182, 314–330. doi:10.1111/j.1469-8137.2009.02773.x.

Rognes, T., Flouri, T., Nichols, B., Quince, C., and Mahé, F. (2016). VSEARCH: a versatile open source tool for metagenomics. PeerJ 4, e2584. doi:10.7717/peerj.2584.

Roscher, C., Schumacher, J., Baade, J., Wilcke, W., Gleixner, G., Weisser, W. W., et al. (2004). The role of biodiversity for element cycling and trophic interactions: an experimental approach in a grassland community. Basic Appl. Ecol. 5, 107–121. doi:10.1078/1439-1791-00216.

Sanders, F. E., and Tinker, P. B. (1973). Phosphate flow into mycorrhizal roots. Pestic. Sci. 4, 385–395. doi:10.1002/ps.2780040316.

Schimel, J. P., and Bennett, J. (2004). Nitrogen mineralization: Challenges of a changing paradigm. Ecology 85, 591–602. doi:10.1890/03-8002.

Shi, S., Nuccio, E. E., Shi, Z. J., He, Z., Zhou, J., and Firestone, M. K. (2016). The interconnected rhizosphere: High network complexity dominates rhizosphere assemblages. Ecol. Lett. 19, 926–936. doi:10.1111/ele.12630.

Stempfhuber, B., Richter-Heitmann, T., Regan, K. M., Kölbl, A., Wüst, P. K., Marhan, S., et al. (2016). Spatial interaction of archaeal ammonia-oxidizers and nitrite-oxidizing bacteria in an unfertilized grassland soil. Front. Microbiol. 6, 1567. doi:10.3389/fmicb.2015.01567.

Strickland, M. S., Osburn, E., Lauber, C., Fierer, N., and Bradford, M. A. (2009). Litter quality is in the eye of the beholder: Initial decomposition rates as a function of inoculum characteristics. Funct. Ecol. 23, 627–636. doi:10.1111/j.1365-2435.2008.01515.x.

Thomson, B. C., Ostle, N. J., McNamara, N. P., Oakley, S., Whiteley, A. S., Bailey, M. J., et al. (2013). Plant soil interactions alter carbon cycling in an upland grassland soil. Front. Microbiol. 4, 253. doi:10.3389/fmicb.2013.00253.

Thuille, A., Laufer, J., Höhl, C., and Gleixner, G. (2015). Carbon quality affects the nitrogen partitioning between plants and soil microorganisms. Soil Biol. Biochem. 81, 266–274. doi:10.1016/j.soilbio.2014.11.024.

Tisserant, E., Malbreil, M., Kuo, A., Kohler, A., Symeonidi, A., Balestrini, R., et al. (2013). Genome of an arbuscular mycorrhizal fungus provides insight into the oldest plant symbiosis. Proc. Natl. Acad. Sci. 110, 20117–20122. doi:10.1073/pnas.1313452110.

Veresoglou, S. D., Chen, B., and Rillig, M. C. (2012). Arbuscular mycorrhiza and soil nitrogen cycling. Soil Biol. Biochem. 46, 53–62. doi:10.1016/J.SOILBIO.2011.11.018.

Vestergård, M., Henry, F., Rangel-Castro, J. I., Michelsen, A., Prosser, J. I., and Christensen, S. (2008). Rhizosphere bacterial community composition responds to arbuscular mycorrhiza, but not to reductions in microbial activity induced by foliar cutting. FEMS Microbiol. Ecol. 64, 78–89. doi:10.1111/j.1574-6941.2008.00447.x.

Wang, Q., Garrity, G. M., Tiedje, J. M., and Cole, J. R. (2007). Naive Bayesian classifier for rapid assignment of rRNA sequences into the new bacterial taxonomy. Appl. Environ. Microbiol. 73, 5261–7. doi:10.1128/AEM.00062-07.

Weisser, W. W., Roscher, C., Meyer, S. T., Ebeling, A., Luo, G., Allan, E., et al. (2017). Biodiversity effects on ecosystem functioning in a 15-year grassland experiment: Patterns, mechanisms, and open questions. Basic Appl. Ecol. 23, 1–73. doi:10.1016/j.baae.2017.06.002.

Whiteside, M. D., Werner, G. D. A., Caldas, V. E. A., van’t Padje, A., Dupin, S. E., Elbers, B., et al. (2019). Mycorrhizal Fungi Respond to Resource Inequality by Moving Phosphorus from Rich to Poor Patches across Networks. Curr. Biol. 29, 2043-2050.e8. doi:10.1016/J.CUB.2019.04.061.

Wickham, H. (2011). ggplot2. Wiley Interdiscip. Rev. Comput. Stat. 3, 180–185. doi:10.1002/wics.147.

Williams, A., Manoharan, L., Rosenstock, N. P., Olsson, P. A., and Hedlund, K. (2017). Long-term agricultural fertilization alters arbuscular mycorrhizal fungal community composition and barley (*Hordeum vulgare*) mycorrhizal carbon and phosphorus exchange. New Phytol. 213, 874–885. doi:10.1111/nph.14196.

Xu, X., Thornton, P. E., and Post, W. M. (2013). A global analysis of soil microbial biomass carbon, nitrogen and phosphorus in terrestrial ecosystems. Glob. Ecol. Biogeogr. 22, 737–749. doi:10.1111/geb.12029.

Zhalnina, K., Louie, K. B., Hao, Z., Mansoori, N., da Rocha, U. N., Shi, S., et al. (2018). Dynamic root exudate chemistry and microbial substrate preferences drive patterns in rhizosphere microbial community assembly. Nat. Microbiol. 3, 470–480. doi:10.1038/s41564-018-0129-3.

Zhang, J., Kobert, K., Flouri, T., and Stamatakis, A. (2014). PEAR: a fast and accurate Illumina Paired-End reAd mergeR. Bioinformatics 30, 614–20. doi:10.1093/bioinformatics/btt593.

Zhu, Q., Riley, W. J., and Tang, J. (2017). A new theory of plant-microbe nutrient competition resolves inconsistencies between observations and model predictions. Ecol. Appl. 27, 875–886. doi:10.1002/eap.1490.

