## Supplementary Information for "Nutrient Source and Mycorrhizal Association jointly alters Soil Microbial Communities that shape Plant-Rhizosphere-Soil Carbon-Nutrient Flows"

**Reference** **14**

### 1 Ingrowth Core

Ingrowth cores are useful as they allow an easy way to implement soil amendments not only in pot experiments but also field experiments. Typically a core has many windows on all sides so as to allow easy access for roots and fungi. The core is also hollow at the top and bottom. The upper hole allows easy venting of respired gases.

Guarding the ingrowth core with meshes enable a control over what grows into them. One of the ways of guarding the core windows (side and bottom) is by applying nylon mesh fabric cut-outs which are often unstable and fall off during manipulation of the core. A more stable method for this is using a mesh bag to encase the entire core (Figure S1).

In this experiment we used two mesh types a 1000 micron and 30 micron mesh. Each mesh was applied as a bag and was given an independent core and pot for implementation. The 30 micron mesh was applied only to the AMF associated plant community to investigate effects of hyphal ingrowth minus the roots. This treatment was subsequently excluded from the experiment as equivalent balancing treatment in the non-mycorrhizal treatment was absent. In terms of carbon nitrogen flow and microbial community composition the 30 micron core was found to be comparable to the 1000 micron core (Table S1) installed in AM plant communities making it a redundant treatment in the experiment as well.

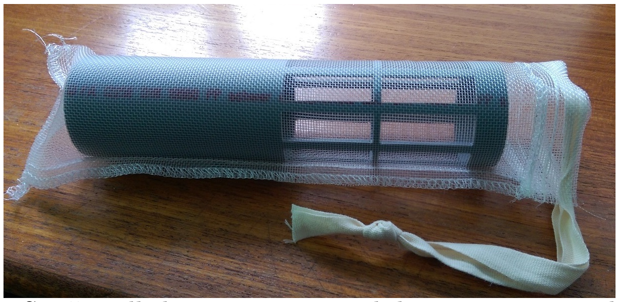

Figure S1: Installed 1000 micron mesh bag on an ingrowth core

Note -

1. No complete root exclusion treatment was carried out where a closed system is used in the experiment.

#### 1.1 Ingrowth Core treatment

Ingrowth cores were removed from the pots two weeks after they were amended with different substrates. The pictures presented in Figure S3 are representative examples of harvested ingrowth cores and offer a visual check of fine root ingrowth into the core after the two week incubation period. This was used as a positive evidence of successful plant influence in combination with delivery of $^{13}C$ into the ingrowth core.

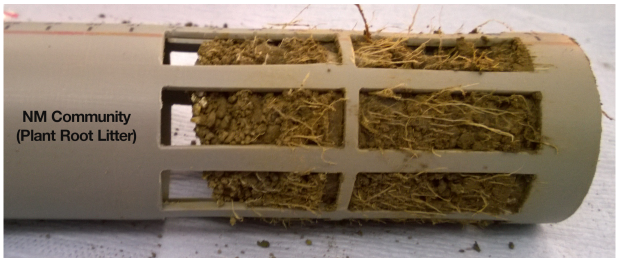

(a) NM Plant Community amended with plant root litter

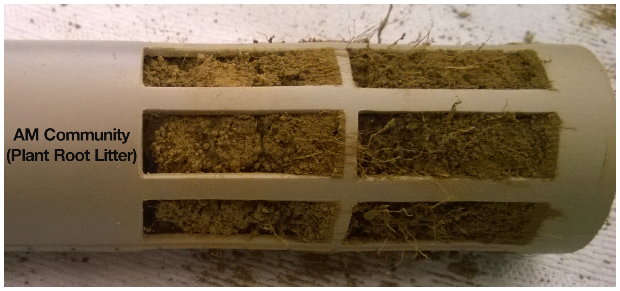

(b) AM plant community amended with plant root litter

Figure S2: Ingrowth Confirmation

#### 1.2 Sieving of Ingrowth core soil

The central position of the ingrowth core and 5 cm diameter resulted in a high degree of root colonization for both plant communities which ensured all of the ingrowth core soil was attached to the root network (Figure S2).The block of soil removed from the ingrowth core was directly deposited on the sieve such that none of the root associated soil is lost. The root attached soil was gently released from the root on to the sieve and alcohol sterilised forceps was used to remove the soil free roots. This strategy of sieving ensured no loss of soil prior to soil amendment and also ensured that we sampled root-free rhizosphere associated soil from the core ensuring all the label measurements on the soil from microbial component of the system.

#### 1.3 Comparison of Mesh types

We isolated the effects of AMF alone by deploying two mesh sizes for the AMF associated plant community. The 30 micron mesh restricted the entry of roots into the ingrowth core soil and thus created hyphosphere which can be defined as soil colonised by AMF hyphae.

Table S1: **Mesh Size Comparison:** Analysis of Variance (ANOVA) comparing different mesh sizes where 1000 micron mesh allows root ingrowth whereas 30 micron mesh restricts entry of roots but allows entry of fungal hypahe. Columns are grouped by measurements.
Replicates: AM community n = 9 and Root excluded n = 9.

| | 15N uptake | | | Respired Carbon | | |
|---|---|---|---|---|---|---|
| Treatment | F.value | P.value | Df | F.value | P.value | Df |
| Mesh Sizes | 0.22 | 0.648 | 1 | 0.16 | 0.695 | 1 |
| Soil Amendment | 0.77 | 0.483 | 2 | 167.65 | < 0.001 (***) | 2 |
| Plant Community x Soil Amendment | 0.12 | 0.891 | 2 | 2.97 | 0.090 | 2 |

Significance is indicated at $p < 0.05$ (*); $p < 0.01$ (**) and $p < 0.001$ (***)

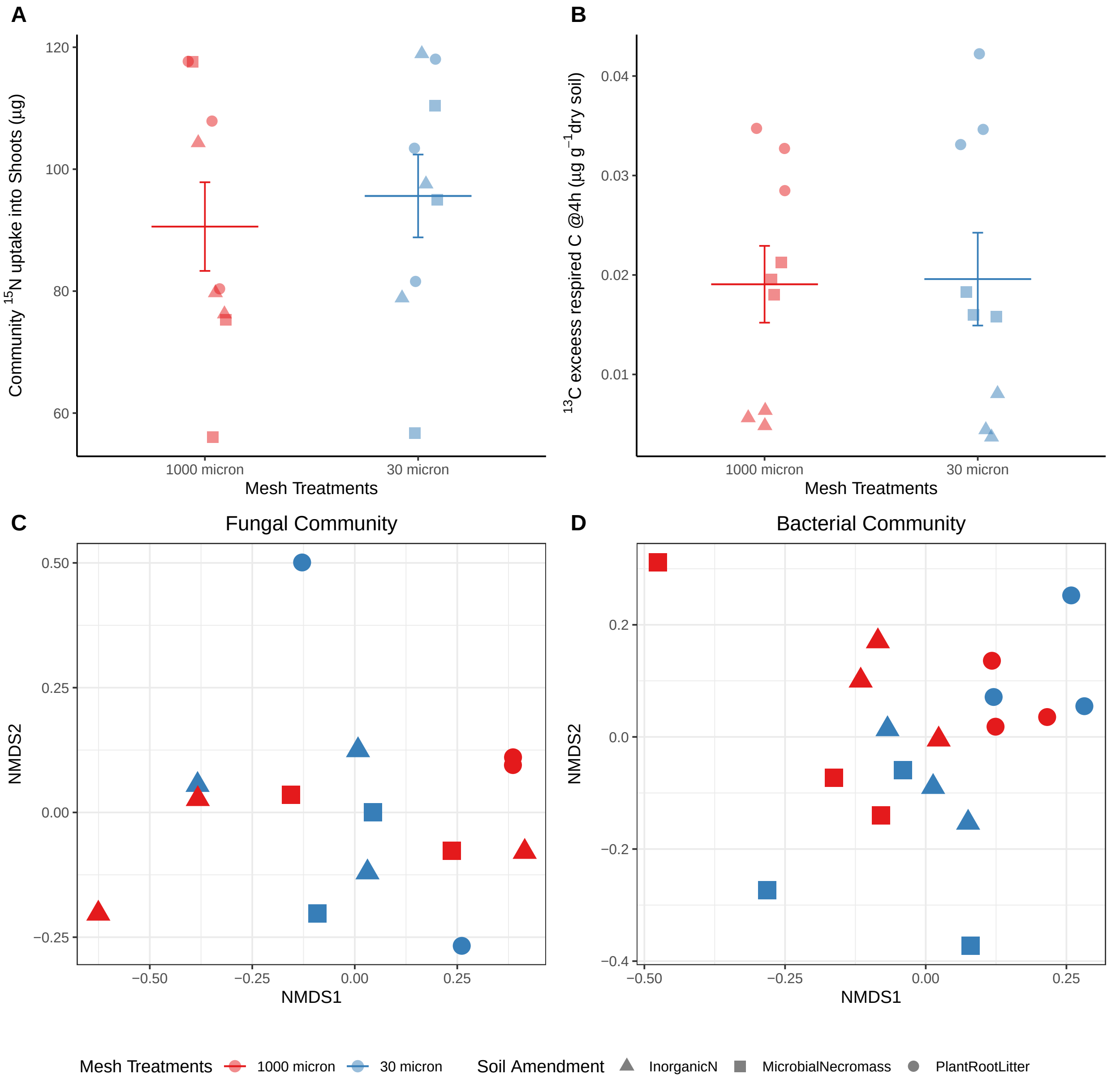

Figure S3: Comparison of two mesh sizes (1000 (Red) and 30 micron (Blue)) on A. Plant N uptake ; B. Plant C uptake into Microbial biomass and C. Plant C respired by microbial community. Microbial community composition in the two meshes were also compared (D: Fungal Community and E: Bacterial Community).

#### 2 Carbon flow in fast pools

##### 2.1 Total Microbial biomass (Fumigation method)

The amount of $^{13}$C excess in fast turnover microbial carbon pool was measured using chloroform fumigation extraction method (Malik et al., 2013). The AM plant community consistently invested much lower amount of freshly photoassimilated carbon into fast turnover microbial biomass pool (Figure S4).

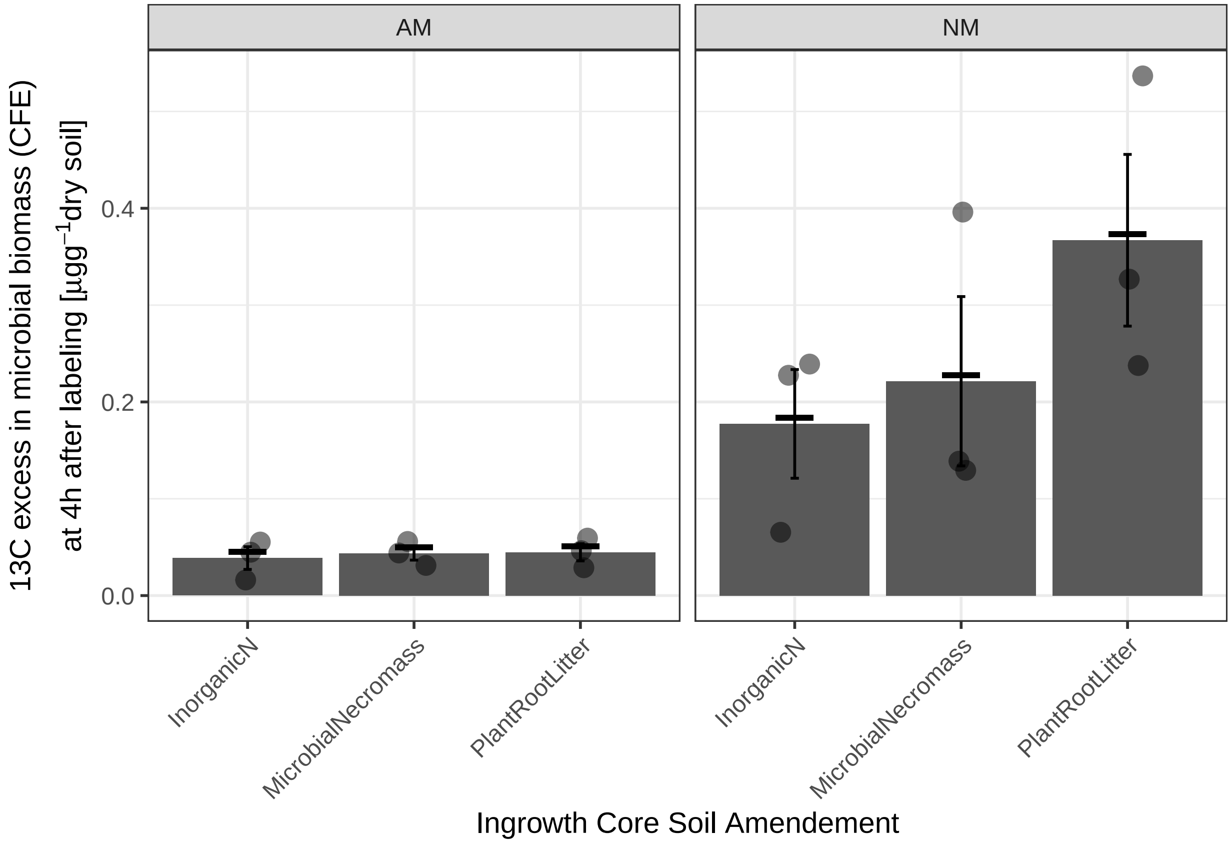

Figure S4: Freshly photoassimilated carbon incroporated into Microbial Biomass at 4h after labelling

#### 2.2 Soil respiration

Approximately 1g of harvested soil at 4h after labelling was incubated for 18 hours in a gas tight vial after which the respired $^{13}CO_2$ content was measured. AM plant community associated microbial communities had a lower $^{13}C$ excess in respired $CO_2$ and consistently respired a lower amount of freshly fixed C compared to NM plant community irrespective of the type of substrate being decomposed.

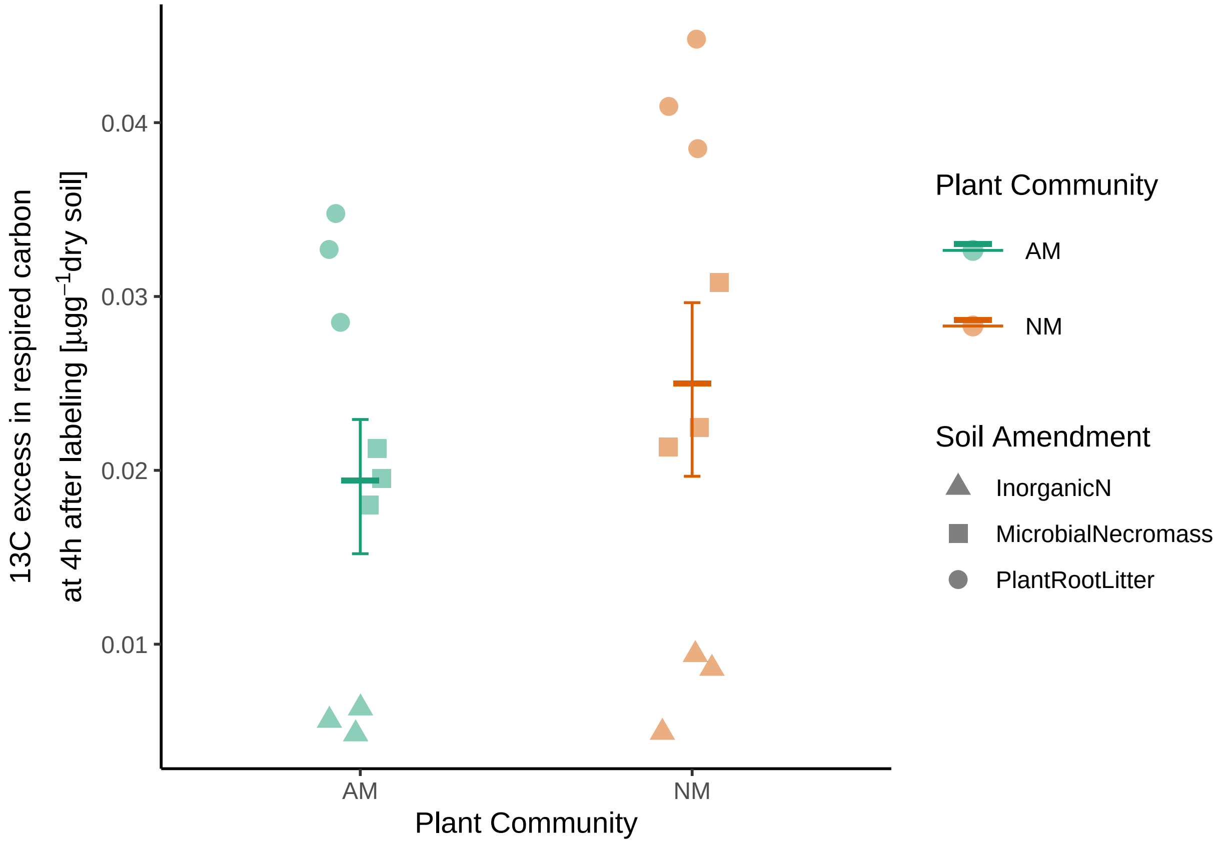

Figure S5: Freshly photoassimilated carbon respired by microbial community in the ingrowth core soil

### 3 Assumpltion for homogeneity of variance (ANOVA)

Table S2: **Test for homogeneity of variances for ANOVA:** Results of Fligner-Killeen test for homogeneity of variance. The substrate amendments were grouped within the plant community types resulting in n=9 per plant community and n=3 for each substrate type grouped under the plant community. Non-significant output satisfies the assumption that variances are homogeneous.

| Variable | Df | Chi.square.value | p.value |
|---|---|---|---|
| Root:Shoot 13C | 5 | 2.76 | 0.735 |
| Microbial Biomass (13C) | 5 | 1.29 | 0.936 |
| Shoot (15N) | 5 | 3.38 | 0.642 |
| Carbon Cost (Total Community) | 5 | 3.73 | 0.590 |
| Carbon Cost (AMF) | 5 | 3.48 | 0.627 |
| Carbon Cost (Fungi) | 5 | 4.52 | 0.478 |
| Carbon Cost (Bacteria) | 5 | 3.13 | 0.680 |

Significance is indicated at $p < 0.05$ (*); $p < 0.01$ (**) and $p < 0.001$ (***)

### 4 Genetic Biomarker Analysis

#### 4.1 Alpha Diversity Indices

As the design of the experiment is hierarchical in the sense that the soil amendments are nested under plant community types being tested, these plots present comparison of alpha diversity indexes for community richness and evenness amongst plant community types within substrate classes used.

##### 4.1.1 Plant Community Comparison

Table S3: **Alpha Diversity indices across plant community:** Comparison of alpha diversity indices for bacterial and fungal communities grouped by AM (n = 9) and NM (n = 8) plant communities.

| | | Mean (SEM) | | ANOVA | | |
|---|---|---|---|---|---|---|
| Microbial.Group | Diversity.Indice | AM.Mean | NM.Mean | Df | F.Value | p.value |
| Bacteria | Observed.OTU | 2198.1 (53.60) | 2174.4 (35.70) | 1 | 0.13 | 0.725 |
| | Shannon.Evenness | 7.2 (0.10) | 7.1 (0.03) | 1 | 0.42 | 0.526 |
| Fungi | Observed.OTU | 455.0 (63.63) | 352.67 (48.81) | 1 | 1.69 | 0.215 |
| | Shannon.Evenness | 5.92 (0.14) | 5.67 (0.16) | 1 | 1.25 | 0.283 |

Significance is indicated at $p < 0.05$ (*); $p < 0.01$ (**) and $p < 0.001$ (***)

##### 4.1.2 Soil Amendment Comparison

Table S4: **Alpha Diversity indices across soil amendments:** Comparison of alpha diversity indices for bacterial and fungal communities grouped by litter type across both plant communities.
Plant root litter (n =5) , Microbial Necromass (n=5) ; Inorganic-N (n=6)

| | | Mean (SEM) | | | ANOVA | | |
|---|---|---|---|---|---|---|---|
| Microbial.Group | Diversity.Indice | Plant.Root.Litter | Microbial.Necromass | Inorganic.N | Df | F.Value | p.value |
| Bacteria | Observed.OTU | 2179 (30) | 2195 (62) | 2187 (83) | 2 | 0.02 | 0.981 |
| | Shannon.Evenness | 7.12 (0.07) | 7.13 (0.05) | 7.14 (0.02) | 2 | 0.05 | 0.948 |
| Fungi | Observed.OTU | 535 (43) | 353 (52) | 319 (71) | 2 | 3.81 | 0.049 (*) |
| | Shannon.Evenness | 6.13 (0.08) | 5.73 (0.11) | 5.54 (0.23) | 2 | 3.16 | 0.076 |

Significance is indicated at $p < 0.05$ (*); $p < 0.01$ (**) and $p < 0.001$ (***)

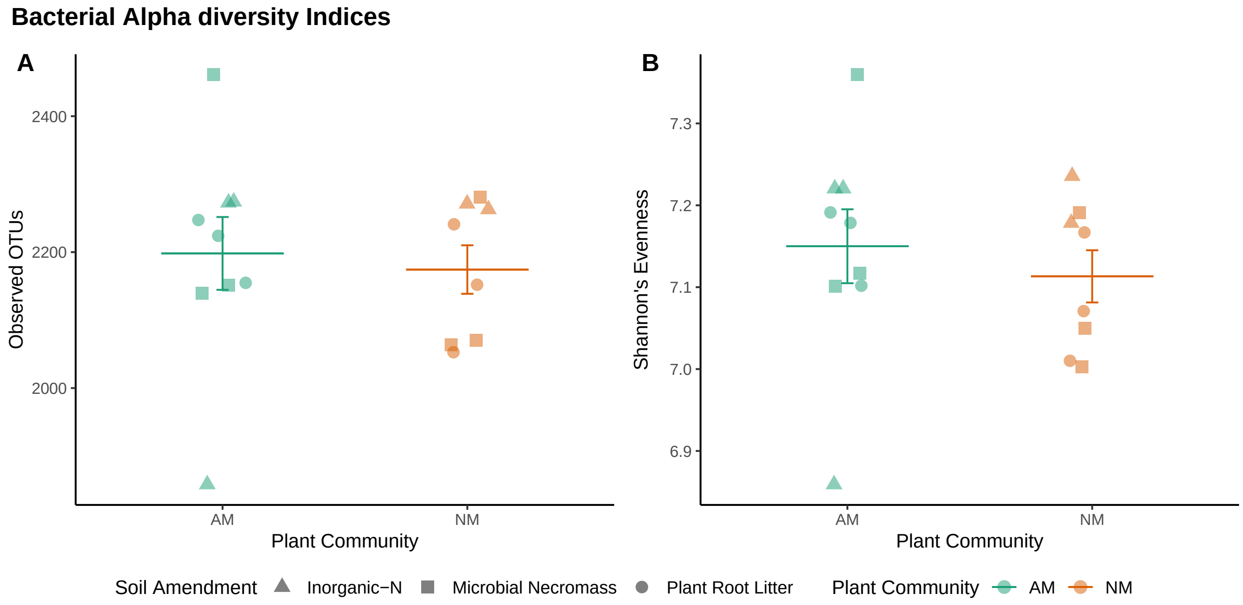

Figure S6: The effect of presence or absence of AMF association on the number of unique bacterial OTUs observed (A) and community evenness (B) when associated soil is amended with biochemically distinct substrates with similar nutrient availability.

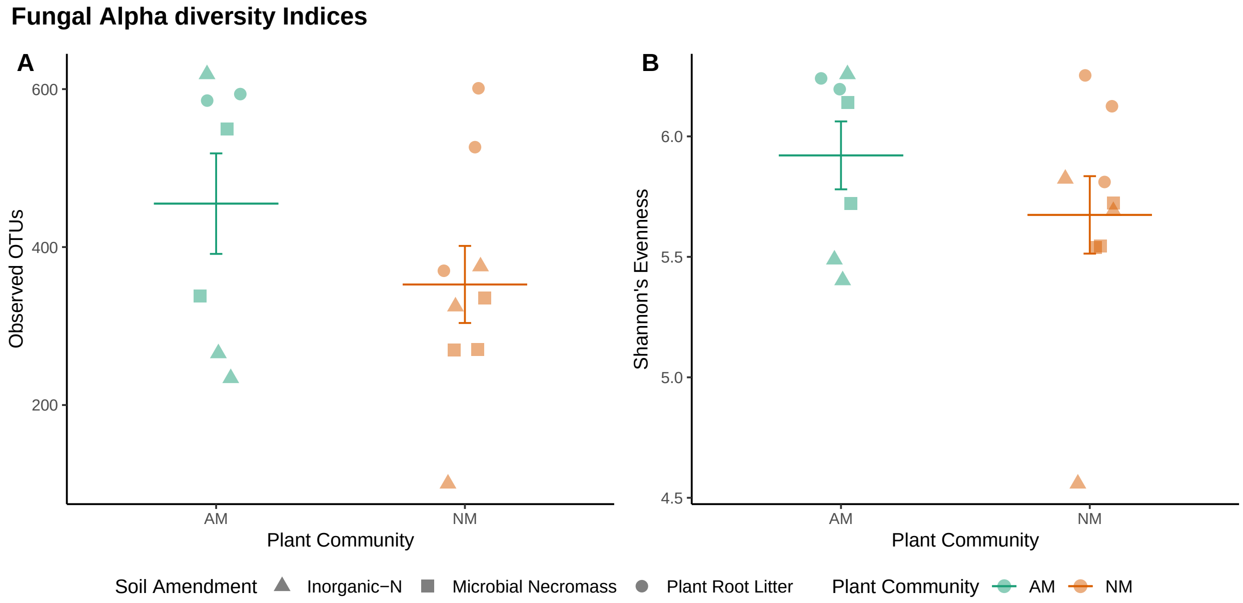

Figure S7: The effect of presence or absence of AMF association on the number of unique Fungal OTUs observed (A) and community evenness (B) when associated soil is amended with biochemically distinct substrates with similar nutrient availability.

#### 4.2 Beta-diversity

##### 4.2.1 Microbial Response to Treatment context

In the experiment two environmental contexts were altered for the microbial communities tested namely plant community type and available substrate resulting from soil amendments where various organic and inorganic substrates were tested. One way analysis of variance was used for each of these context. In case of bacteria relative abundance of individual OTUs were grouped by specific phylogenetic assignment to ascertain group response to experimental variables i.e. if the microbial group is influenced by plant community context or by alteration to substrates they feed upon. For fungi relative abundance of individual OTUs were grouped by guild assignments and subjected to similar one way analysis of variance as the bacterial groups.

Table S5: **Bacterial Response to Plant Community:** Mean (SEM) relative abundance of each bacterial group for each plant community type irrespective of soil amendment. F value and significance represents one way anova results to determine responsiveness to plant community context.
Replicates: AM community n = 9 and NM community n = 8.

| | Mean (SEM) | | ANOVA | |
|---|---|---|---|---|
| Taxa | Rel.Abundance.in.AM | Rel.Abundance.in.NM | F.value | p.value |
| *Acidobacteria* | 0.241 (0.010) | 0.307 (0.021) | 8.21 | 0.012 (*) |
| *Actinobacteria* | 0.076 (0.003) | 0.064 (0.003) | 6.24 | 0.025 (*) |
| *Bacteroidetes* | 0.117 (0.005) | 0.107 (0.011) | 0.79 | 0.389 |
| *Choloroflexi* | 0.027 (0.001) | 0.027 (0.002) | 0.02 | 0.903 |
| *Gemmamonadetes* | 0.022 (0.002) | 0.018 (0.002) | 2.73 | 0.120 |
| *Nitrospirae* | 0.016 (0.002) | 0.017 (0.002) | 0.32 | 0.578 |
| *Proteobacteria* | 0.358 (0.011) | 0.312 (0.012) | 8.38 | 0.011 (*) |
| *Verrucomicrobia* | 0.103 (0.007) | 0.106 (0.006) | 0.11 | 0.741 |
| Others | 0.038 (0.002) | 0.041( 0.001) | 1.42 | 0.253 |
| *Alphaproteobateria* | 0.139 (0.005) | 0.124 (0.006) | 3.62 | 0.076 |
| *Betaproteobacteria* | 0.076 (0.003) | 0.063 (0.003) | 8.36 | 0.011 (*) |
| *Deltaproteobacteria* | 0.073 (0.005) | 0.066 (0.002) | 2.09 | 0.168 |
| *Gammaproteobacteria* | 0.070 (0.004) | 0.059 (0.004) | 3.89 | 0.067 |

Significance is indicated at p < 0.05 (*); p < 0.01 (**) and p < 0.001 (***)

Table S6: **Fungal Guild Response to Plant Community:** Mean (SEM) relative abundance of each fungal guild for each plant community type irrespective of soil amendment. F value and significance represents one way anova results to determine responsiveness to plant community context.
Replicates: AM community n = 7 and NM community n = 9.

| | Mean (SEM) | | ANOVA | |
|---|---|---|---|---|
| Taxa | Rel.Abundance.in.AM | Rel.Abundance.in.NM | F.value | Sign |
| Saprotroph | 0.345 (0.023) | 0.493 (0.015) | 32.310 | < 0.001 (***) |
| AMF | 0.370 (0.035) | 0.120 (0.009) | 58.370 | < 0.001 (***) |
| Ectomycorrhizal | 0.013 (0.002) | 0.012 (0.002) | 0.078 | 0.784 |
| Endophyte | 0.233 (0.023) | 0.324 (0.022) | 8.26 | 0.012 (*) |
| Others | 0.040 (0.004) | 0.051 (0.010) | 0.919 | 0.354 |

Significance is indicated at p < 0.05 (*); p < 0.01 (**) and p < 0.001 (***)

Table S7: **Bacterial Response to Soil Amendment:** Mean (SEM) relative abundance of each bacterial group for each soil amendment type irrespective of plant community. F value and significance represents one way anova results to determine responsiveness to soil amendment.
Replicates: Plant Root Litter n = 6 ; Microbial Necromass n = 6 and Inorganic-N n = 5.

| | Mean (SEM) | | | ANOVA | |
|---|---|---|---|---|---|
| Taxa | Plant.Root.Litter | Microbial.Necromass | Inorganic.N | F.value | p.value |
| *Acidobacteria* | 0.299 (0.029) | 0.279 (0.018) | 0.223 (0.005) | 2.718 | 0.101 |
| *Actinobacteria* | 0.068 (0.003) | 0.070 (0.006) | 0.068 (0.005) | 0.351 | 0.710 |
| *Bacteroidetes* | 0.096 (0.009) | 0.114 (0.008) | 0.128 (0.009) | 3.244 | 0.070 |
| *Choloroflexi* | 0.029 (0.002) | 0.027 (0.001) | 0.026 (0.002) | 0.987 | 0.397 |
| *Gemmamonadetes* | 0.024 (0.002) | 0.015 (0.002) | 0.021 (0.001) | 7.866 | 0.005 (**) |
| *Nitrospirae* | 0.021 (0.001) | 0.014 (0.001) | 0.014 (0.001) | 21.880 | <0.001 (***) |
| *Proteobacteria* | 0.325 (0.018) | 0.331 (0.019) | 0.359 (0.009) | 1.102 | 0.359 |
| *Verrucomicrobia* | 0.089 (0.002) | 0.111 (0.008) | 0.113 (0.010) | 3.296 | 0.067 |
| Others | 0.045 (0.002) | 0.037 (0.002) | 0.036 (0.003) | 5.787 | 0.015 (*) |
| *Alphaproteobateria* | 0.121 (0.005) | 0.136 (0.008) | 0.140 (0.007) | 2.008 | 0.171 |
| *Betaproteobacteria* | 0.066 (0.004) | 0.071 (0.006) | 0.072 (0.003) | 0.398 | 0.679 |
| *Deltaproteobacteria* | 0.076 (0.003) | 0.059 (0.003) | 0.075 (0.004) | 7.253 | 0.007 (**) |
| *Gammaproteobacteria* | 0.061 (0.006) | 0.064 (0.004) | 0.072 (0.004) | 1.086 | 0.364 |

Significance is indicated at $p < 0.05$ (*); $p < 0.01$ (**) and $p < 0.001$ (***)

Table S8: **Fungal Guild Response to Soil Amendment:** Mean (SEM) relative abundance of each fungal guild for each soil amendment type irrespective of plant community. F value and significance represents one way anova results to determine responsiveness to soil amendment.
Replicates: Plant Root Litter n = 5 ; Microbial Necromass n = 5 and Inorganic-N n = 6.

| | Mean (SEM) | | | ANOVA | |
|---|---|---|---|---|---|
| Taxa | Plant.Root.Litter | Microbial.Necromass | Inorganic.N | F.value | p.value |
| Saprotroph | 0.465 (0.0422) | 0.442 (0.022) | 0.387 (0.045) | 1.103 | 0.361 |
| AMF | 0.224 (0.067) | 0.191 (0.045) | 0.265 (0.073) | 0.341 | 0.717 |
| Ectomycorrhizal | 0.014 (0.003) | 0.011 (0.003) | 0.012 (0.003) | 0.136 | 0.874 |
| Endophyte | 0.231 (0.020) | 0.314 (0.034) | 0.304 (0.035) | 2.023 | 0.172 |
| Others | 0.067 (0.010) | 0.042 (0.005) | 0.032 (0.009) | 4.534 | 0.032 (*) |

Significance is indicated at $p < 0.05$ (*); $p < 0.01$ (**) and $p < 0.001$ (***)

##### 4.2.2 Assumption Testing for Permanova

Betadisper was used to calculate homogeneity of group dispersions (Oksanen et al., 2018).

Table S9: **Test for homogeneity of variance assumption:** Multivariate homogeneity of group dispersions (variances) using betadisper was tested for fungal and bacterial community for the entire experimental setup. F value and p value reported represents results of analysis of varience (ANOVA) on the betadisper output.

| | | ANOVA | | |
|---|---|---|---|---|
| Community | Factor | d.f | F.value | p.value |
| Fungi | Plant Community | 1 | 0.9728 | 0.399 |
| | Soil Amendment | 2 | 1.9751 | 0.143 |
| | Plant Community : Soil Amendment | 5 | 1.5002 | 0.271 |
| Bacteria | Plant Community | 1 | 0.0340 | 0.856 |
| | Soil Amendment | 2 | 0.4083 | 0.692 |
| | Plant Community : Soil Amendment | 5 | 2.3314 | 0.099 |

Significance is indicated at $p < 0.05$ (*); $p < 0.01$ (**) and $p < 0.001$ (***)

##### 4.2.3 Permutational Multivariate Analysis of Variance (PERMANOVA)

Adonis function was used to calculate PERMANOVA output (Oksanen et al., 2018).

Table S10: Permutational Multivariate Analyses of Variance (PERMANOVA) of soil fungal community matrix from ingrowth core soil decomposing various litter types in presence or absence of AMF association.

| Community | Exp.Variable | Df | F.value | R.squared | p.value |
|---|---|---|---|---|---|
| Fungal community | Plant Community | 1 | 1.87 | 0.11 | 0.013 (*) |
| | Soil Amendment | 2 | 1.99 | 0.23 | 0.0003 (**) |
| | Plant Community x Soil Ammendment | 2 | 0.90 | 0.10 | 0.677 |

Significance is indicated at $p < 0.05$ (*); $p < 0.01$ (**) and $p < 0.001$ (***)

Table S11: Permutational Multivariate Analyses of Variance (PERMANOVA) of soil bacterial community matrix from ingrowth core soil decomposing various litter types in presence or absence of AMF association .

| Community | Exp.Variable | Df | F.value | R.squared | p.value |
|---|---|---|---|---|---|
| Bacterial community | Plant Community | 1 | 1.24 | 0.07 | 0.019 (*) |
| | Soil Amendment | 2 | 1.50 | 0.17 | 0.0001 (***) |
| | Plant Community x Soil Ammendment | 2 | 1.08 | 0.12 | 0.129 |

Significance is indicated at $p < 0.05$ (*); $p < 0.01$ (**) and $p < 0.001$ (***)

### 5 Data Availability

#### 5.1 Bacterial 16S rRNA gene data

**Download URLS and metadata for Bacterial 16S rRNA gene amplicon data**

Table S12: Bacterial amplicon data at NCBI SRA

| SRA.Exp.Accession | SRA.Run.number | Sample.ID | Plant.Community | Soil.Amendment | Download.URL |
|---|---|---|---|---|---|
| SRX3647783 | SRR6671091 | D1 | NM | Plant Root Litter | click here |
| SRX3647784 | SRR6671090 | D2 | NM | Plant Root Litter | click here |
| SRX3647785 | SRR6671089 | D3 | NM | Plant Root Litter | click here |
| SRX3647781 | SRR6671093 | D7 | AM | Plant Root Litter | click here |
| SRX3647782 | SRR6671092 | D8 | AM | Plant Root Litter | click here |
| SRX3647777 | SRR6671097 | D9 | AM | Plant Root Litter | click here |
| SRX3647778 | SRR6671096 | D37 | NM | Microbial Necromass | click here |
| SRX3647795 | SRR6671079 | D38 | NM | Microbial Necromass | click here |
| SRX3647796 | SRR6671078 | D39 | NM | Microbial Necromass | click here |
| SRX3647792 | SRR6671082 | D43 | AM | Microbial Necromass | click here |
| SRX3647789 | SRR6671085 | D44 | AM | Microbial Necromass | click here |
| SRX3647790 | SRR6671084 | D45 | AM | Microbial Necromass | click here |
| SRX3647787 | SRR6671087 | D73 | NM | Inorganic N | click here |
| SRX3647788 | SRR6671086 | D75 | NM | Inorganic N | click here |
| SRX3647771 | SRR6671103 | D79 | AM | Inorganic N | click here |
| SRX3647776 | SRR6671098 | D80 | AM | Inorganic N | click here |
| SRX3647775 | SRR6671099 | D81 | AM | Inorganic N | click here |

#### 5.2 Fungal ITS data

**Download URLS and metadata for Fungal ITS region cDNA based amplicon data**

Table S13: Fungal amplicon data at NCBI SRA

| SRA.Exp.Accession | SRA.Run.number | Sample.ID | Plant.Community | Soil.Amendment | Download.URL |
|---|---|---|---|---|---|
| SRX4378626 | SRR7508173 | R1 | NM | Plant Root Litter | click here |
| SRX4378627 | SRR7508172 | R2 | NM | Plant Root Litter | click here |
| SRX4378624 | SRR7508175 | R3 | NM | Plant Root Litter | click here |
| SRX4378623 | SRR7508176 | R8 | AM | Plant Root Litter | click here |
| SRX4378620 | SRR7508179 | R9 | AM | Plant Root Litter | click here |
| SRX4378621 | SRR7508178 | R37 | NM | Microbial Necromass | click here |
| SRX4378628 | SRR7508171 | R38 | NM | Microbial Necromass | click here |
| SRX4378629 | SRR7508170 | R39 | NM | Microbial Necromass | click here |
| SRX4378643 | SRR7508156 | R43 | AM | Microbial Necromass | click here |
| SRX4378636 | SRR7508163 | R45 | AM | Microbial Necromass | click here |
| SRX4378637 | SRR7508162 | R73 | NM | Inorganic-N | click here |
| SRX4378638 | SRR7508161 | R74 | NM | Inorganic-N | click here |
| SRX4378639 | SRR7508160 | R75 | NM | Inorganic-N | click here |
| SRX4378630 | SRR7508169 | R79 | AM | Inorganic-N | click here |
| SRX4378633 | SRR7508166 | R80 | AM | Inorganic-N | click here |
| SRR7508167 | SRX4378632 | R81 | AM | Inorganic-N | click here |

### Reference

Malik, A., Blagodatskaya, E., & Gleixner, G. (2013). Soil microbial carbon turnover decreases with increasing molecular size. *Soil Biol. Biochem.*, *62*, 115–118. https://doi.org/10.1016/j.soilbio.2013.02.022

Oksanen, J., Blanchet, F. G., Friendly, M., Kindt, R., Legendre, P., McGlinn, D., Minchin, P. R., O'Hara, R. B., Simpson, G. L., Solymos, P., Stevens, H. H., Szoecs, E., & Wagner, H. (2018). *vegan: Community Ecology Package*. https://cran.r-project.org/package=vegan
